# An RNA-based feed-forward mechanism ensures motor switching in oskar mRNA transport

**DOI:** 10.1101/2021.04.24.441269

**Authors:** Imre Gáspár, Ly Jane Phea, Mark A. McClintock, Simone Heber, Simon L. Bullock, Anne Ephrussi

## Abstract

Regulated recruitment and activity of motor proteins is essential for intracellular transport of cargoes, including messenger ribonucleoprotein complexes (RNPs). Here we show that orchestration of *oskar* RNP transport in the *Drosophila* germline relies on the interplay of two double-stranded RNA binding proteins, Staufen and the dynein adaptor Egalitarian (Egl). We find that Staufen antagonizes Egl-mediated transport of *oskar* mRNA by dynein both *in vitro* and *in vivo*. Following delivery of nurse cell-synthesized *oskar* mRNA into the oocyte by dynein, recruitment of Staufen to the RNPs results in dissociation of Egl and a switch to kinesin-1-mediated translocation of the mRNA to its final destination at the posterior pole of the oocyte. We additionally show that Egl associates with *staufen* (*stau)* mRNA in the nurse cells, mediating its enrichment and translation in the ooplasm. Our observations identify a novel feed-forward mechanism, whereby dynein-dependent accumulation of *stau* mRNA, and thus protein, in the oocyte enables motor switching on *oskar* RNPs by downregulating dynein activity.

## INTRODUCTION

Proper execution of the genetic program relies on regulatory mechanisms that act at multiple levels of gene expression, from transcription to post-translational events. In many cell types, including neurons, early embryos and oocytes, a key process that regulates genetic output at the subcellular level is RNA localization, which is often driven by cytoskeletal motors (St Johnston, 2005; Gaspar, 2011; Marchand *et al.* 2012; Glock *et al.* 2017; Mofatteh and Bullock, 2017). The kinesin-1 and cytoplasmic dynein-1 (dynein) motors, which move RNPs towards the plus end and minus end of microtubules, respectively, play a central role in mRNA trafficking in several systems (Gaspar, 2011; Mofatteh and Bullock, 2017).

A paradigm for the study of microtubule-based RNP transport is *oskar* mRNA, the protein product of which induces abdomen and germline formation in the embryo (Ephrussi and Lehmann, 1992). *oskar* mRNA is produced in the nurse cells of the *Drosophila* germline syncytium in the early stages of oogenesis and transported by dynein into the developing oocyte where microtubule minus ends are enriched (Ephrussi *et al.* 1991; Januschke *et al.*, 2002; Clark *et al.* 2007; Sanghavi *et al.*, 2013; Jambor *et al.*, 2014; Vazquez-Pianzola *et al.*, 2017). mRNA trafficking from the nurse cells to the oocyte is dependent on the Egl protein, which binds specialized stem-loop structures in localizing mRNAs and links them together with its coiled-coil binding partner Bicaudal-D (BicD) to dynein and the dynein activating complex dynactin (Navarro *et al.*, 2004; Dienstbier *et al.*, 2009; Dix *et al.*, 2013). *In vitro* reconstitution studies have shown that Egl and BicD also play a key role in switching on dynein motility (McClintock *et al.*, 2018; Sladewski *et al.*, 2018).

During mid-oogenesis, when the microtubule cytoskeleton is reorganized, most of Egl’s mRNA targets localize to the anterior of the oocyte, where microtubule minus ends are now concentrated (Gaspar, 2011; Lasko, 2012). In contrast, *oskar* mRNA is transported to the posterior pole, where the microtubule plus ends are focused (Parton *et al.*, 2011); (Sanghavi *et al.*, 2013), by the Kinesin-1 heavy chain (Khc) (Brendza *et al.*, 2000; Palacios and St Johnston, 2002; Zimyanin *et al.*, 2008; Williams *et al.*, 2014). Khc is associated with nascent *oskar* RNPs in the nurse cell cytoplasm (Gáspár *et al.*, 2017), raising the question of how the activity of the motor is coordinated with that of dynein during delivery of *oskar* mRNA from the nurse cells to the oocyte posterior.

Another double-stranded RNA binding protein (dsRBP), Staufen, is necessary for the Khc-mediated localization of *oskar* to the posterior pole (Ephrussi *et al.* 1991; St Johnston *et al*. 1991; Januschke *et al.*, 2002; Sanghavi *et al.*, 2013; Jambor *et al.*, 2014; Vazquez-Pianzola *et al.*, 2017; St Johnston *et al.*, 1992; Zimyanin *et al.*, 2008), as well as for production of Oskar protein at this site (St Johnston *et al.* 1991; Ephrussi and Lehmann, 1992; Schuldt *et al.*, 1998; Micklem *et al.*, 2000). Staufen is also involved in trafficking of other mRNAs, such as *bicoid* in the oocyte and *prospero* in dividing neuroblasts (Ferrandon *et al.*, 1994; Schuldt *et al.*, 1998). Mammalian orthologues of *Drosophila* Staufen, mStau1 and mStau2, play roles in the bidirectional transport of *CaMKII*α and *Rgs4* mRNAs in dendrites (Heraud-Farlow *et al.*, 2013; Bauer *et al.*, 2019) and *Xenopus* Staufen is a component of *Vg1* transport RNPs in the oocyte (Yoon and Mowry, 2004). These observations indicate that Staufen’s function as a regulator of RNP transport is evolutionarily conserved. However, it is unclear how Staufen proteins orchestrate mRNA trafficking processes.

Here, we show that Staufen antagonizes dynein-based transport of *oskar* RNPs within the *Drosophila* oocyte and thus favors kinesin-1-driven translocation of the mRNA to the posterior pole. Recombinant Staufen can inhibit dynein-based transport of *oskar* mRNPs that are reconstituted *in vitro* with purified components, showing that it is a direct regulator of the transport machinery. Staufen displaces Egl from *oskar* RNPs after they have reached the oocyte. Staufen does not, however, perturb dynein and dynactin association with the RNPs, revealing that proteins in addition to Egl contribute to motor linkage in this system. These data suggest that Staufen interferes with dynein activation by promoting dissociation of Egl from RNPs. We also provide evidence that Staufen’s inhibition of dynein is restricted to the oocyte by Egl- and dynein-mediated concentration of nurse cell-synthesized *stau* mRNA within the ooplasm. Thus, Egl delivers its own negative regulator into the oocyte, constituting a feed-forward regulatory mechanism that spatially and temporally controls dynein activity and thus *oskar* mRNA localization during oogenesis.

## RESULTS

### Staufen suppresses minus end-directed transport of *oskar* RNPs in ooplasmic extracts

*oskar* mRNA is localized at the posterior pole of the oocyte during mid-oogenesis (Ephrussi *et al.* 1991; Kim-Ha *et al.* 1991). In *stau* mutant oocytes, enrichment of *oskar* mRNA at the posterior pole is greatly reduced, and a considerable amount of *oskar* remains at the anterior margin (Fig 1A’; (Ephrussi *et al.* 1991; St Johnston *et al.* 1991). To assess the effect of *stau* depletion on the balance of *oskar* directionality on microtubules, we evaluated the movement of individual MCP-GFP-labeled *oskar-MS2* RNPs in control and *stau* RNAi ooplasmic extracts by TIRF microscopy (Gaspar and Ephrussi, 2017; Gáspár *et al.*, 2017). This analysis revealed that the normal ~ 2:1 dominance of microtubule plus end-versus minus end-directed *oskar* RNP runs (Gaspar and Ephrussi, 2017; Gáspár *et al.*, 2017) was lost, such that there was no overt bias in the directionality of motility (Fig 1B).

**Figure 1:**
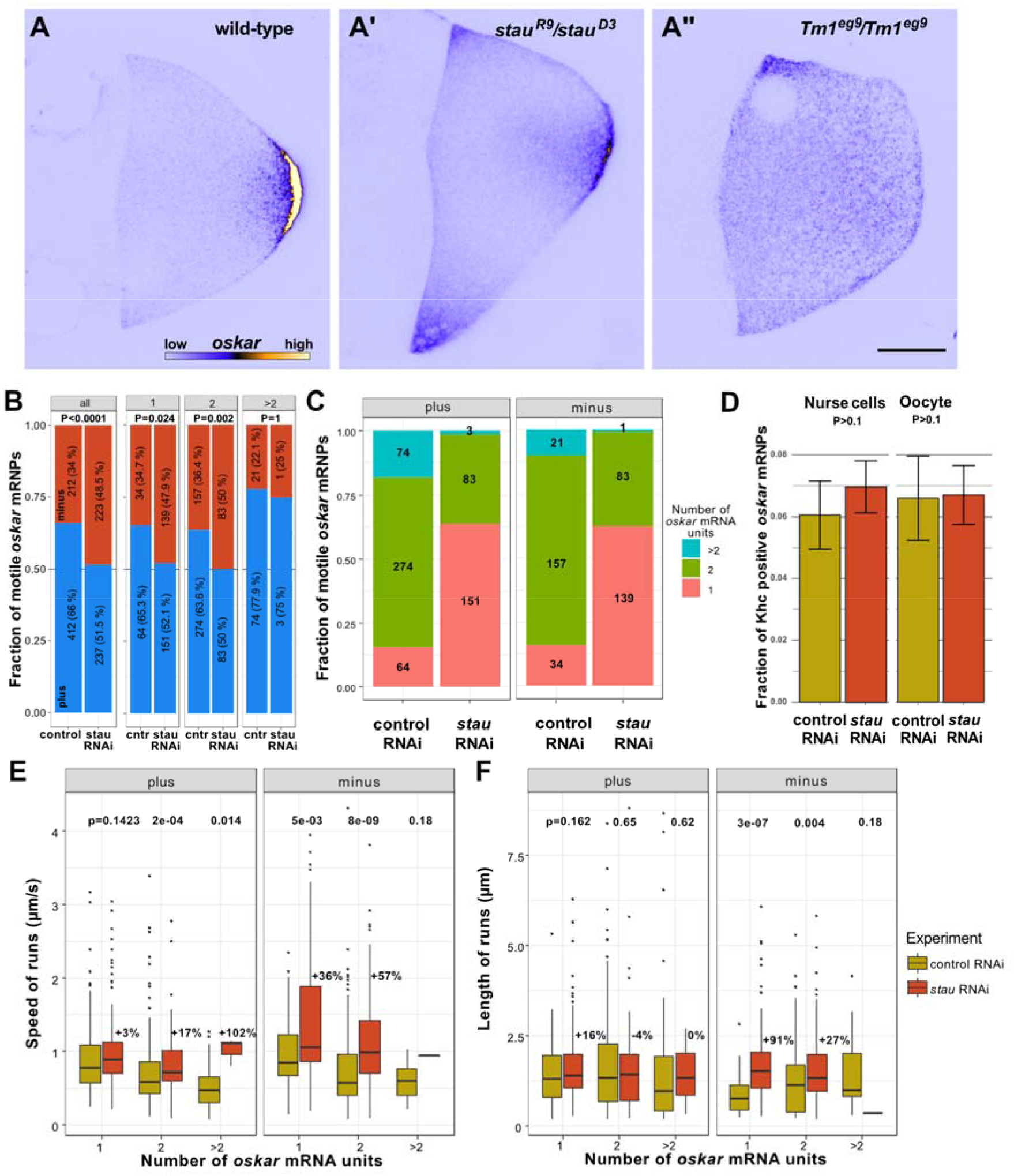
*oskar* RNA localization and transport are impaired in *stau* mutants. (A-A”) Endogenous *oskar* mRNA localization (shown in blue (low intensity)/yellow (high intensity)) in wild-type (A), *stau* null (A’) and *Tm1-I/C* null stage 9 oocytes (A”) (anterior to the left and posterior to the right). Scale bar represents 20 μm. In *stau* null oocytes, there is a pronounced anterior accumulation of *oskar* in addition to the normal posterior localization; in Tm1-I/C mutant oocytes, *oskar* RNA is predominantly enriched at the anterior cortex. (B) Polarity distribution of *oskar*-MS2 RNP runs in control RNAi and *stau* RNAi ooplasmic extracts. Numbers indicate the number of *oskar* RNP runs measured for all motile *oskar* RNPs and for RNPs categorized by relative RNA content (Video S3 and S4). (C) Relative mRNA content of motile *oskar-MS2* RNPs in control RNAi and *stau* RNAi ooplasmic extracts. Pink, green and blue indicate the fraction of *oskar* RNPs with one, two, or more relative RNA units, respectively (see Methods). Numbers show the number of *oskar* RNPs in that category. (D) Association of Khc and *oskar* RNPs *in situ* in the nurse cells and the oocyte of control RNAi versus *stau* RNAi samples. (E-F) Speed (E) and run length (F) of motile *oskar-MS2* RNPs towards the plus or minus ends of microtubules in control RNAi and *stau* RNAi ooplasmic extracts. RNPs are stratified by relative RNA content (D). Numbers above the boxplots (E and F) indicate the differences in mean travel distance between *stau* and control RNAi extracts. In *stau* RNAi extracts there were too few motile *oskar-MS2* RNPs with >2 RNAs to establish a statistical trend (C,E-F). P-values of Fisher exact (B) or pairwise Mann-Whitney U-tests (D-F) are shown.

While analyzing the motility, we noticed that the relative GFP intensity of the motile fraction of *oskar* RNPs in the *stau* RNAi extracts was lower than in the control RNAi condition, indicating fewer mRNA molecules on average. This observation might explain the paucity of *oskar* mRNA transport detected in live *stau* mutant oocytes when using the less sensitive confocal live-cell imaging technique (compare Videos S1-S4; (Zimyanin *et al.*, 2008)). However, loss of Staufen does not seem to affect the overall RNA content of *oskar* RNPs in the oocyte as revealed by single-molecule fluorescent *in situ* hybridization (smFISH) (Fig S5B), indicating a selective effect on the size of motile RNPs.

Since differences in cargo size and/or composition could confound the analysis, we compared motility of *oskar* RNPs of similar sizes by stratifying the GFP intensity of RNPs into ‘relative’ RNA units (see Methods and Fig 1C). This approach confirmed that motile *oskar* RNPs contained fewer RNA molecules on average in the *stau* RNAi condition (Fig 1C) and additionally showed that the loss of plus-end dominance could be observed for motile RNPs with 1 or 2 relative units of *oskar* RNA, which make up the majority of the motile population (Fig 1B). These data show that Staufen is needed for the plus-end dominance that localizes *oskar* to the posterior pole.

The distribution of *oskar* mRNA and the directionality of RNP transport in *stau* mutants differ from what is observed in other mutants affecting *oskar* localization: in *Tm1^gs^* (Fig 1A” and (Gáspár *et al.*, 2017) and *Khc^null^* (Brendza *et al.*, 2000; Zimyanin *et al.*, 2008; Williams *et al.*, 2014; Gáspár *et al.*, 2017) mutants, *oskar* mRNA does not localize at the posterior of the oocyte, and minus-end-directed transport predominates as a failure of Khc mediated posterior-ward transport (Gáspár *et al.*, 2017). To explore the basis of the directionality defect in *stau* mutants, we quantified the colocalization in egg chambers of functional, endogenously tagged fluorescent Khc molecules and *oskar* mRNA detected by smFISH. This method was previously used to show that Tm1 mutations reduce the association of *oskar* RNPs with the motor (Gáspár *et al.*, 2017). When Staufen was depleted, there was no reduction in Khc colocalization with *oskar* mRNA in either the nurse cells or the oocyte (Fig 1D), indicating that the observed loss of plus-end-directed bias (Fig 1B) is not due to reduced association of Khc with *oskar* RNPs.

To assess whether there is a change in motor activity when Staufen is disrupted, we analyzed the speed and run lengths of GFP-labeled *oskar-MS2* RNPs towards both microtubule ends in *stau* RNAi and control ooplasmic extracts. The velocity of *oskar* RNP movement appears to be influenced by their RNA content, as larger RNPs moved more slowly in either direction (Fig 1E). When Staufen was depleted, there was a substantial and statistically significant increase in minus-end-directed run lengths and velocities of RNPs containing 1 or 2 relative units of *oskar* RNA (Fig 1D). In contrast, features of plus-end-directed motion were generally not affected in the *stau* RNAi condition with the exception of a modest, but significant, increase in plus end velocity for RNPs with 2 *oskar* RNA units (Fig 1E). These data indicate that the loss of plus-end-directed dominance of *oskar* RNPs when Staufen is depleted is predominantly associated with hyperactivity of the minus end-directed dynein transport machinery. Previous work with gain-of-function mutations in the dynein activator BicD showed that increased dynein activity leads to ectopic anterior accumulation of *oskar* RNA (Navarro *et al.*, 2004; Liu *et al.*, 2013), as is observed in *stau* mutant oocytes (Fig 1A’,(St Johnston *et al.*, 1991)). Altogether, these findings point to a role of Staufen in suppression of minus end-directed transport of *oskar* RNA by dynein.

### Staufen antagonizes dynein-mediated *oskar* mRNA transport by purified components

To test if Staufen has a direct inhibitory effect on minus-end directed *oskar* RNP transport, we made use of an *in vitro* motility assay that involves reconstitution of a minimal RNP assembled from purified Egl, BicD, dynactin, dynein and *in vitro* transcribed RNA (McClintock *et al.*, 2018). Motility of this RNP along immobilized, fluorescent microtubules can be visualized by TIRF microscopy using fluorophores on the mRNA and dynein.

In these experiments we used a ~529 nt fragment of the *oskar* 3’UTR that is sufficient for nurse cell-to-oocyte transport (Jambor *et al.*, 2014). In the presence of Egl, BicD and dynactin, *oskar* mRNA frequently underwent long-distance transport in association with dynein (Fig. 2A and Fig S2A). No transport of *oskar* was detected when Egl and BicD were omitted from the mix (Fig. S2A); in this condition, association of *oskar* mRNA with microtubules or processive movement of microtubule-bound dynein complexes was almost never observed (Fig. S2A-C). These observations corroborate the conclusion that Egl and BicD link mRNAs to dynein-dynactin and activate movement of the motor along microtubules (Dienstbier *et al.*, 2009; McClintock *et al.*, 2018; Sladewski *et al.*, 2018).

**Figure 2.**
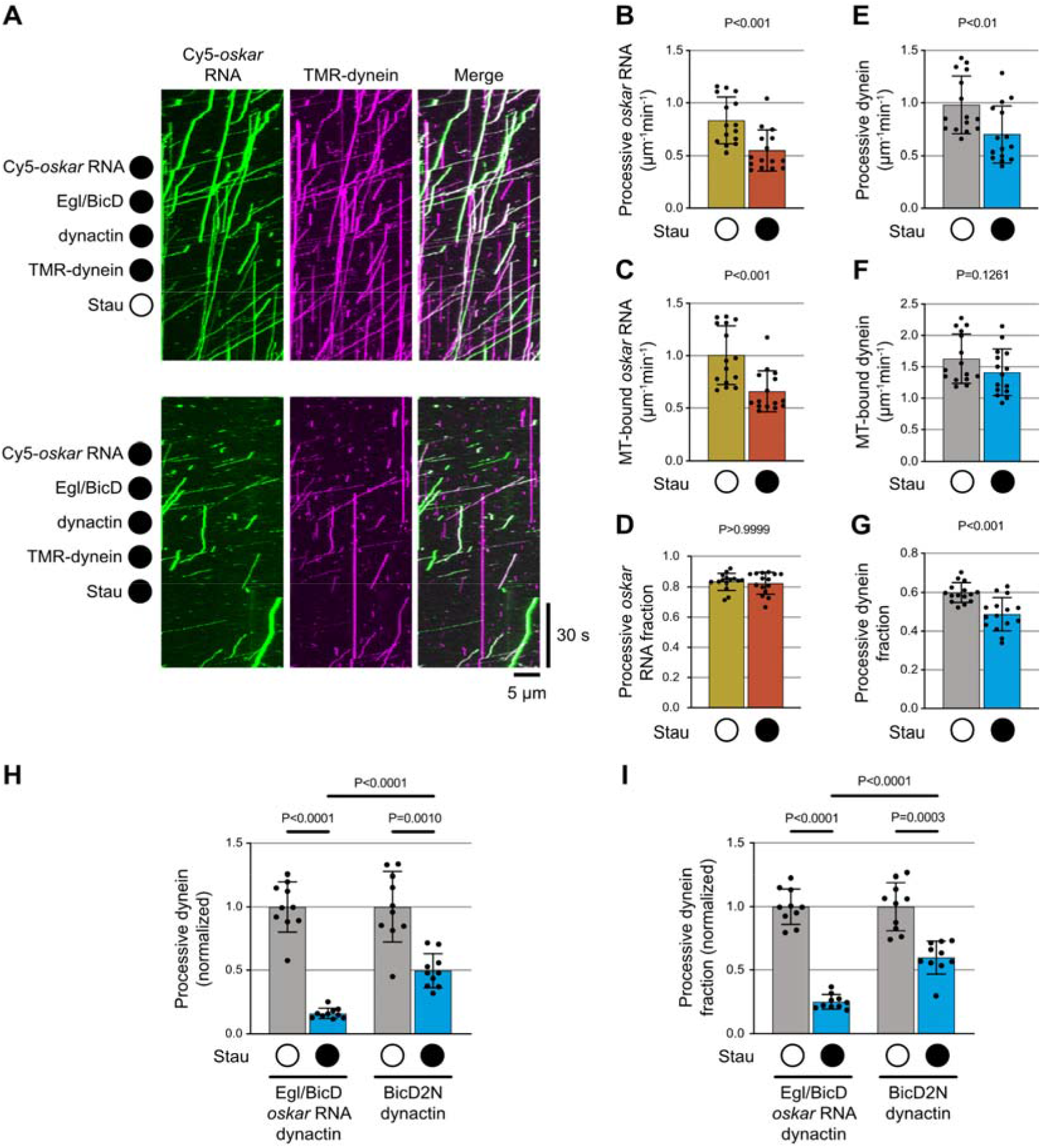
Staufen impairs *oskar* mRNA transport and dynein activation *in vitro*. (A) Kymographs (time-distance plots) exemplifying the behavior of *oskar* RNA and dynein in the presence (filled circle) and absence (open circle) of Staufen; in both conditions, Egl, BicD and dynactin are also present but not fluorescently labeled. Minus end is to the left and plus end to the right. (B-G) Quantification of motile properties of *oskar* RNA (B-D) and dynein (E-G) in the conditions shown in A. Charts show frequency of processive movements (B, E), total number of microtubule (MT) binding events (C, F) and fraction of microtubule-binding events that result in processive motility (D, G). (H-I) Quantification of effect of Staufen on number of processive dynein complexes and fraction of dynein-microtubule binding events that result in processive motility for motor activated by dynactin, *oskar* RNA, and Egl/BicD (H) vs motor activated by dynactin and BicD2N (I). In H and I, values for conditions with Staufen were normalized to the corresponding condition without Staufen to obtain relative metrics. Plots show the mean ± standard deviation (SD) of values from individual microtubules (represented by black circles) derived from analysis of 586-1341 single RNA particles or 1247-2207 single dynein particles per condition. Statistical significance and P-values were determined with Mann-Whitney tests (B-G) or with Brown-Forsythe and Welch ANOVA tests (H and I).

Addition of recombinant Staufen to the Egl, BicD, dynein and dynactin assembly mix significantly reduced the number of *oskar* mRNA transport events (Fig. 2A and B). This effect was associated with fewer mRNAs being recruited to the microtubules (Fig. 2C). However, those mRNAs that interacted with the microtubule were just as likely to undergo long-distance transport as those incubated without Staufen (Fig. 2D). The presence of Staufen also reduced the number of processive dynein complexes and the proportion of microtubule-associated dyneins that underwent processive movement (Fig. 2E-G). We conclude from these experiments that Staufen directly inhibits transport of *oskar* by Egl, BicD, dynactin and dynein and that this effect is associated with impaired activation of motor movement.

Dynein movement during mRNA transport is switched on by the N-terminal coiled-coil region of BicD, which recruits dynactin (and thereby its activating functions) to the motor complex (Dienstbier *et al.*, 2009; McKenney *et al.*, 2010, 2014; Schlager *et al.*, 2014; Hoogenraad and Akhmanova, 2016). These events are triggered by docking of RNA-bound Egl to the C-terminal region of BicD, which frees the N-terminal region from an autoinhibitory interaction (McClintock *et al.*, 2018; Sladewski *et al.*, 2018). To better resolve how dynein activity is impaired by Staufen, we assessed its effect on dynein activation by the N-terminal coiled-coil of the mouse BicD orthologue BicD2 (BicD2N), which is constitutively active in the absence of cargo (Hoogenraad *et al.*, 2003; Dienstbier *et al.*, 2009; McKenney *et al.*, 2010, 2014; Schlager *et al.*, 2014; Hoogenraad and Akhmanova, 2016). In the presence of dynactin and BicD2N, Staufen partially reduced the number of processive dynein movements and the proportion of processive dyneins (Fig. 2H, I and Fig. S3). However, Staufen’s inhibitory effect on the motor was proportionally much stronger in the presence of *oskar*, Egl, full-length BicD and dynactin (Fig. 2H, I and Fig S3). These data suggest that Staufen inhibits both the activation of dynein-dynactin motility by BicD proteins, as well as stimulation of this event by Egl and RNA.

### A balance of Staufen and Egl is required for *oskar* mRNA localization *in vivo*

We next explored the interplay between Egl and Staufen function *in vivo* by manipulating the dosage of these proteins and determining the effects on *oskar* mRNA localization in mid-oogenesis. We evaluated mRNA distributions qualitatively (Fig 3) and confirmed our conclusions through an established, unbiased statistical averaging method (Ghosh *et al.*, 2012; Gaspar *et al.*, 2014) (Fig S4). Similar to the situation in *stau* mutant egg-chambers (Fig 3A, B and Figs S1A, B, E, S4A and B), UAS/Gal4-mediated overexpression of Egl in the germline resulted in a strong *oskar* signal at the anterior of the oocyte cortex, and a weaker enrichment of the mRNA at the posterior (Fig 3C and S4C). This finding corroborates recent observations in Egl-overexpressing egg-chambers by Mohr *et al.* (Mohr *et al.*, 2021).

**Figure 3:**
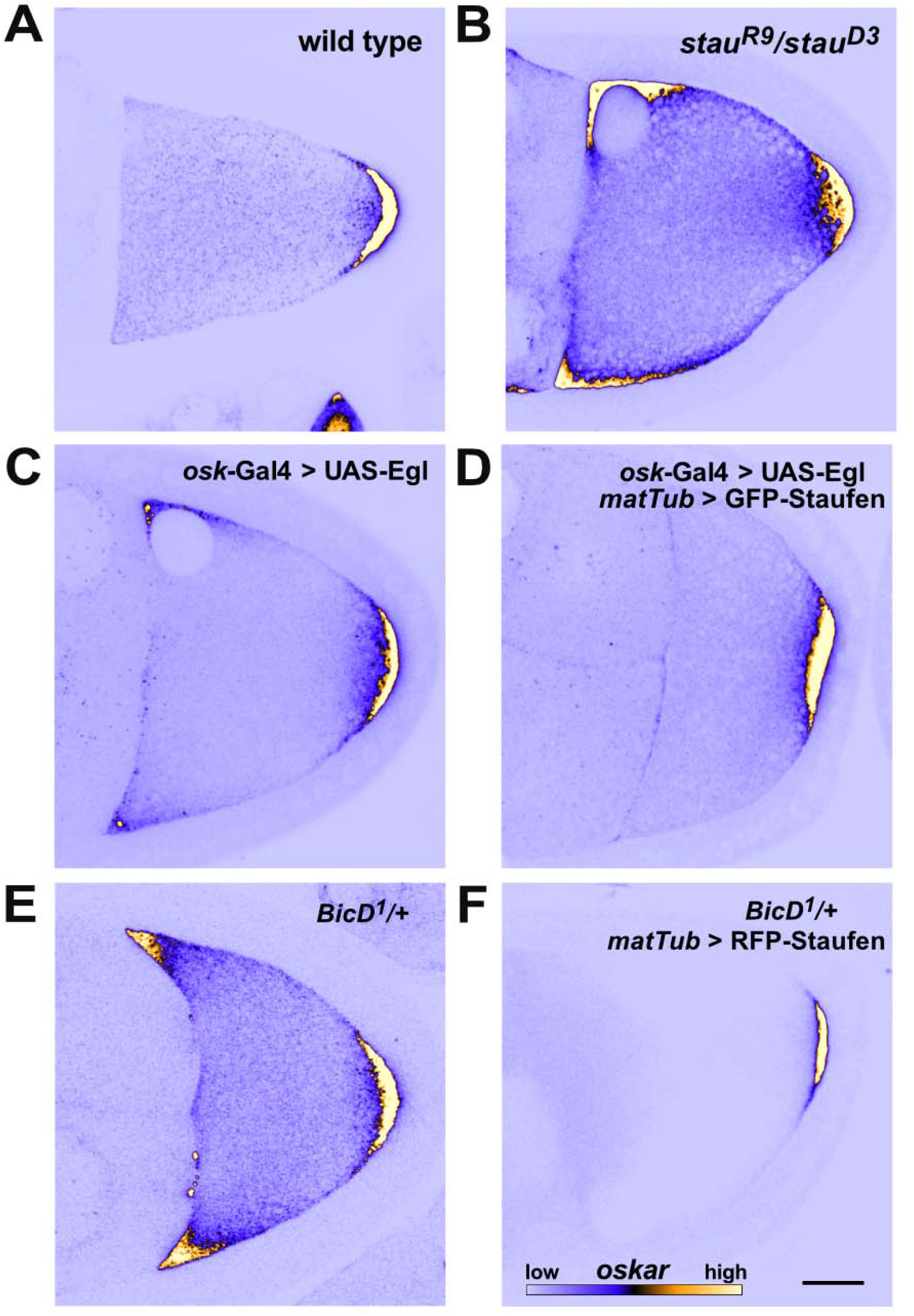
Mutual interference of Staufen and Egl during *oskar* mRNA localization. (A-F) Representative micrographs of the distribution of endogenous *oskar* mRNA (detected by smFISH; blue - low intensity, yellow - high intensity, see scale in F) in stage 9 oocytes (anterior to the left, posterior to the right). Ectopic anterior accumulation of *oskar* mRNA in oocytes lacking Staufen protein (B; see also Fig 1A’), overexpressing Egl (C) or heterozygous for one copy of the dominant, hyperactive *BicD^1^* allele (E). Anterior accumulation of *oskar* is not observed in wild-type oocytes (A), or upon Staufen overexpression in Egl-overexpressing oocytes (D) or *BicD^1^/+* oocytes (F). Scale bar represents 20 μm.

Altogether, these observations suggest that Staufen and Egl antagonize one another during *oskar* mRNA localization in mid-oogenesis. To further test this idea, we overexpressed both GFP-Staufen (using the maternal tubulin promoter (Micklem *et al.* 2000)) and Egl in the germline and examined *oskar* localization. While GFP-Staufen overexpression in this manner had no major effect on the localization of *oskar* (Fig S1F-I and (Heber *et al.*, 2019)), it suppressed the ectopic anterior localization of *oskar* mRNA caused by elevated Egl levels (Fig 3C, D and Fig S4C, D). This observation is consistent with mutual antagonism between Egl and Staufen. Further supporting this notion, removal of a functional copy of *egl* partially suppressed *oskar* mislocalization at the anterior of egg-chambers with reduced Staufen levels (*stau* RNAi, *egl^1^/+*; Fig S3C-E). Furthermore, overexpression of Staufen suppressed the ectopic anterior localization of *oskar* mRNA caused by two different dominant *BicD* alleles (Fig 3E, F and Fig S4E-H). At least one of these mutations (*BicD^1^*) appears to suppress the autoinhibited state of BicD in which the dynein and Egl binding sites are folded back on one another (Liu *et al.*, 2013). Antagonism between BicD and Staufen is further supported by the observation that reduced *stau* gene dosage enhances embryonic lethality associated with the dominant *BicD* alleles (Navarro *et al.*, 2004). Collectively, these genetic interaction experiments indicate a mutually antagonistic relationship between Staufen and Egl, as well as the latter protein’s binding partner, BicD, during the localization of *oskar* mRNA in mid-oogenesis.

### Staufen dissociates Egl from *oskar* RNPs in the oocyte

To shed light on how Staufen and Egl influence each other *in vivo*, we examined the association of each protein with *oskar* mRNA using smFISH in combination with transgenically-expressed, fluorescently-tagged proteins (Gáspár *et al.*, 2017; Heber *et al.*, 2019).

Distribution of transgenic Staufen-GFP, expressed by the *stau* promoter at close to endogenous levels (Fig S1I) resembles that of the endogenous protein (St Johnston *et al.* 1991), and this molecule is functional during *oskar* localization (Fig S1D, E and G). In the oocyte, Staufen-GFP and *oskar* mRNA frequently overlapped. In the nurse cells, there was little overlap between Staufen-GFP and *oskar*, as previously reported (Little *et al.*, 2015), and only low levels of Staufen-GFP signal were observed, regardless of the developmental stage of the egg-chamber (Fig 4A, A’ and B). In the oocyte, the fraction of Staufen-associated *oskar* RNPs increased as a function of their *oskar* mRNA content (which was determined by calibration of smFISH signals (see Materials and Methods and (Little *et al.*, 2015)); Fig S4A). This increase was substantially more pronounced during stage 9, when there is a strong enrichment of *oskar* at the oocyte posterior, than in earlier stages of oogenesis (Fig 4B). During stage 9, not only the likelihood of Staufen association with *oskar* RNPs increased but also the relative levels of Staufen per RNP (Fig 4B’). We previously observed the same behavior for overexpressed GFP-Staufen and mammalian GFP-Stau2 (Heber *et al.*, 2019). Notably, however, no or greatly reduced scaling of the Staufen signal with *oskar* mRNA content of the RNPs was observed in the oocyte prior to the onset of *oskar* localization to the posterior (Fig 4B’). This is likely due to limited availability of the protein, as suggested by the relatively low fluorescent signal in early stage Staufen-GFP oocytes (Fig S5G). Collectively, our analysis shows that Staufen preferentially associates with *oskar* RNA in the oocyte and that this association is largely governed by the RNA content of the RNPs and the enrichment of the pool of Staufen protein in the ooplasm.

**Figure 4:**
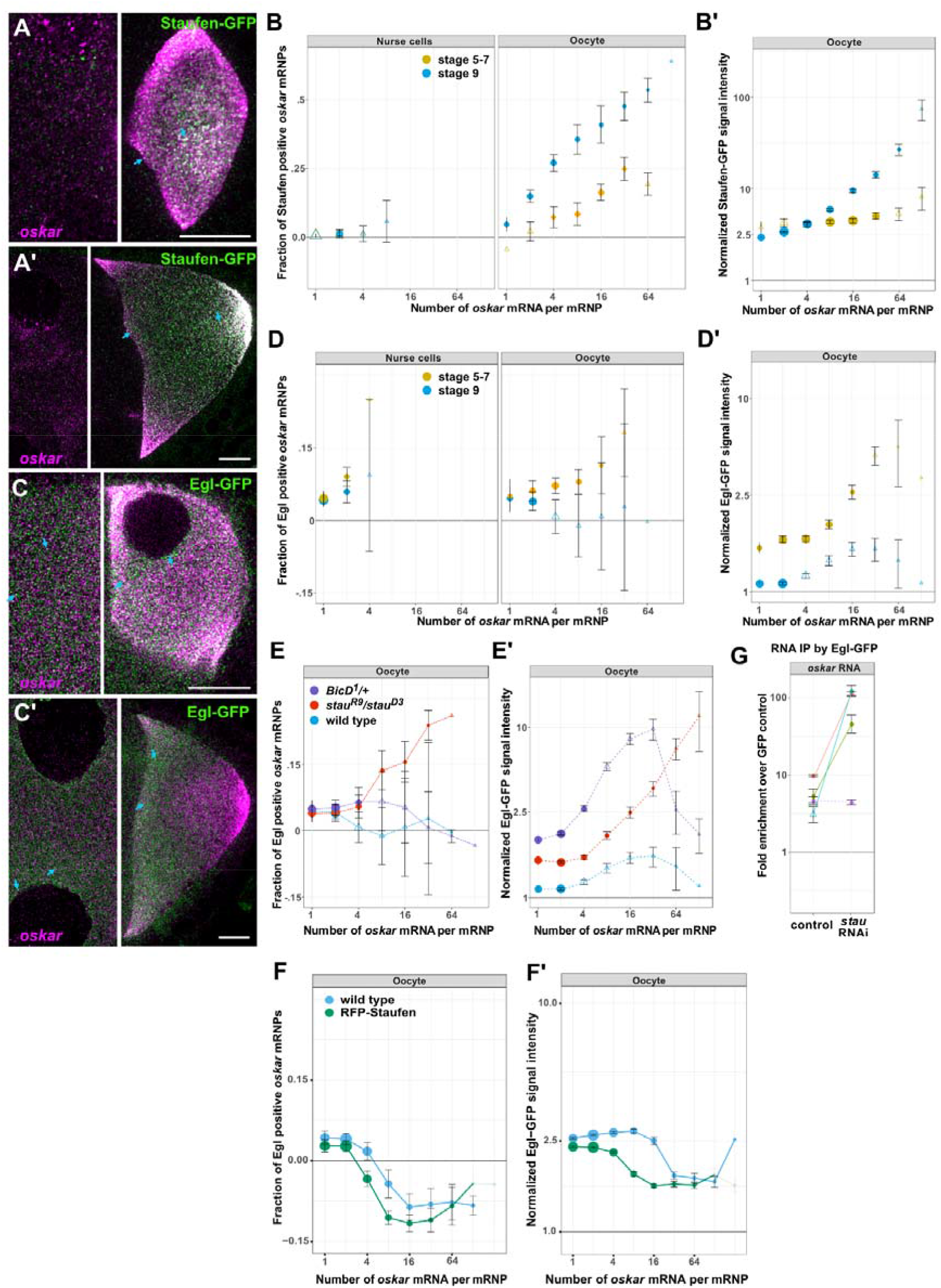
Staufen interferes with association of Egl with *oskar* RNPs in the oocyte. (A-A’) Staufen-GFP (green) distribution in the nurse cell cytoplasm (left-hand images) and oocyte (right-hand images) before (stage 6-7, A) and during (stage 8-9, A’) the posterior localization of endogenous *oskar* mRNA (magenta) in the oocyte of the same egg-chamber. (B-B’) Quantification of Staufen-GFP association with *oskar* RNPs in nurse cell cytoplasm and ooplasm at the indicated stages as a function of RNA copy number. (C-C’) Distribution of Egl-GFP (green) in the nurse cell cytoplasm (left-hand images) and oocyte (right-hand images) before (stage 7, C) and during (stage 9, C’) the posterior localization of endogenous *oskar* mRNA (magenta) in the oocyte of the same egg-chamber. In A, A’, C and C’, nurse cell images are shown with different brightness/contrast settings for better visualization of the signals. Cyan arrows point to examples of colocalization. Scale bars represent 10 μm. (D-D’) Quantification of association of Egl-GFP with *oskar* RNPs in nurse cell cytoplasm and ooplasm at the indicated stages. (E-F’) Association of Egl-GFP with *oskar* RNPs in the stage 9 oocyte in the indicated genotypes. In A-B’ endogenous Staufen was absent (*stau^R9^/stau^D3^* background), whereas in C-F’ endogenous, unlabeled Egl was present. Error bars represent 95% confidence intervals in B, B’, D-F’. The size of the circles is proportional to the relative abundance of each category of *oskar* RNPs within the overall population. Triangles indicate that the fraction of GFP-positive *oskar* RNPs is not significantly different from zero (p>0.01, B, B’, D-F’). In B’, D’, E’ and F’, GFP protein intensities on *oskar* RNPs in the oocytes are normalized to GFP signal intensities in the corresponding nurse cells (G) Fold-enrichment of *oskar* mRNA precipitated from UV crosslinked Egl-GFP ovarian extracts relative to the GFP control in control and *stau* RNAi. Different colors represent different experiments. Solid (p<0.05) or dashed lines (p>0.05) connect paired data (pairwise Student’s t-test was used to determine significance). Empty triangles indicate non-significant enrichment (p>0.05) of *oskar* relative to the GFP control. Error bars show SDn.

During the same stages of oogenesis, Egl-GFP (which was mildly overexpressed with the α-tubulin84B promoter (Fig S5I, (Dienstbier *et al.*, 2009)) was abundant in both the nurse cells and the oocyte (Fig 4C and C’). This pattern matches the distribution of endogenous Egl protein, as revealed by immunostaining (Mach and Lehmann, 1997). Colocalization analysis showed that, in contrast to Staufen-GFP, Egl-GFP was already associated with *oskar* RNPs in the nurse cells (Fig 4D) and maintained its association with *oskar* in the oocyte until the onset of posterior localization. However, from stage 9 onward, we detected no significant colocalization between Egl and RNPs containing four or more copies of *oskar* mRNA (Fig 4D). The amount of Egl signal associated with the RNPs increased proportionately with *oskar* mRNA content during early oogenesis but reduced by about two-fold and became independent of *oskar* mRNA copy number during stage 9 (Fig S5D’). As this loss of Egl from *oskar* RNPs was an unexpected finding, we wondered if it was caused by the dilution of Egl-GFP by endogenous, untagged Egl. However, this was not the case as the reduction of Egl association with *oskar* RNPs was robust to changes in the dosage of the Egl-GFP transgene or endogenous unlabeled Egl and was also observed when no unlabeled Egl (*egl^1^/egl^2^* rescued by the Egl-GFP transgene) was present in the oocyte (Fig S5F’). Collectively, these results indicate that whereas Staufen levels increase on *oskar* RNPs as posterior-ward movement in the oocyte is initiated, there is a concomitant decrease in Egl levels on these complexes.

The opposing trends of Staufen and Egl association with *oskar* RNPs in the nurse cells and the oocyte (Fig 4B,B’ and D,D’), together with the increasing concentrations of Staufen in the ooplasm prior to the onset of *oskar* localization (Fig S5G), are consistent with Staufen inhibiting Egl association with *oskar* RNPs in the oocyte. To test this notion, we analyzed the association of Egl-GFP with *oskar* RNPs in *stau* mutants. When Staufen protein was absent or greatly reduced, Egl remained associated with *oskar* RNPs in stage 9 oocytes, and the relative amount of Egl signal that colocalized with the RNPs increased with *oskar* mRNA content (Fig 4E, E’ and Fig S5 E, E’). Conversely, when Staufen was mildly overexpressed using an RFP-Staufen transgene (Zimyanin *et al.*, 2008), the amount of Egl associated with *oskar* RNPs in the oocyte (in particular with those containing 4+ copies of RNA) decreased (Fig 4F, F’).

To test biochemically whether Staufen impairs the association of Egl with *oskar* RNPs *in vivo*, we performed UV crosslinking and GFP immunoprecipitation followed by quantitative RT-PCR on egg-chambers co-expressing Egl-GFP and either *stau* RNAi or a control RNAi in the germline (Fig 4G, Fig S5H). Substantially more *oskar* mRNA was co-immunoprecipitated with Egl-GFP from extracts of egg-chambers expressing *stau* RNAi compared to the control (Fig 4G). We conclude from these experiments that Staufen antagonizes the association of Egl with *oskar* mRNA *in vivo*.

### Aberrant anterior localization of *oskar* mRNA correlates with increased Egl association

As described above, BicD is a direct binding partner of Egl, and contacts the dynein and dynactin complexes (Dienstbier *et al.*, 2009; Liu *et al.*, 2013; Vazquez-Pianzola *et al.*, 2017; McClintock *et al.*, 2018; Sladewski *et al.*, 2018). To gain further insight into how the association of Egl influences trafficking of *oskar* mRNA, we examined the effect of the two gain-of-function *BicD* mutations in more detail. Both alleles caused an increase in the association of Egl-GFP with mid- to large-sized (4-16 RNA copies) *oskar* RNPs in stage 9 oocytes when compared to the wild-type control (Fig 4E, E’ and Fig S5F, F’). Although the magnitude of the effect differed between the two *BicD* alleles, in both cases the amount of Egl signal on the RNPs scaled with *oskar* RNA copy number up to 32 copies, in contrast with the low amount of Egl detected on *oskar* RNPs in wild-type stage 9 egg-chambers (Fig 4E’ and Fig S5F’). Thus, there is a correlation between the ability of the BicD gain-of-function alleles to misdirect *oskar* RNPs to the anterior of the oocyte and their stimulation of Egl association with these structures in the oocyte cytoplasm.

We reasoned that the anterior localization of *oskar* mRNA in the oocyte observed when untagged Egl was strongly overexpressed (Fig 3C and S4C) might be due to an aberrant, concentration-dependent retention of the protein on the RNPs. To test this idea, we introduced the moderately overexpressed Egl-GFP transgene (Dienstbier *et al.*, 2009) into egg-chambers overexpressing untagged Egl under UAS-Gal4 control (Fig. S5I). We observed numerous large RNP granules containing high amounts of Egl-GFP (which presumably also contain unlabelled Egl) and *oskar* mRNA (Fig 5A and A’) in the nurse cells and at the anterior cortex of the oocyte. Such RNPs were seldom detected in the absence of UAS/Gal4-mediated Egl overexpression (Fig 5A and E). Quantitative analysis revealed that the relative Egl-GFP signal scaled with *oskar* RNA content in the RNPs when untagged Egl was overexpressed (Fig 5C’). The effect was similar to that observed in the absence of Staufen, as well as in egg-chambers with *BicD* gain-of-function alleles (Fig 4E’ and S4F’). In contrast, we found no significant change in Staufen association with *oskar* RNPs in stage 9 oocytes strongly overexpressing Egl (Fig 5D and D’). This finding indicates that Egl does not overtly interfere with the loading of Staufen on *oskar* RNPs. However, substantially more high copy number *oskar* RNPs (containing 17+ copies of *oskar*) were localized at the anterior pole when Egl was strongly overexpressed than in the wild type (Fig 5E), consistent with our earlier finding that Egl antagonizes Staufen’s function in promoting posterior localization of *oskar* mRNA.

**Figure 5:**
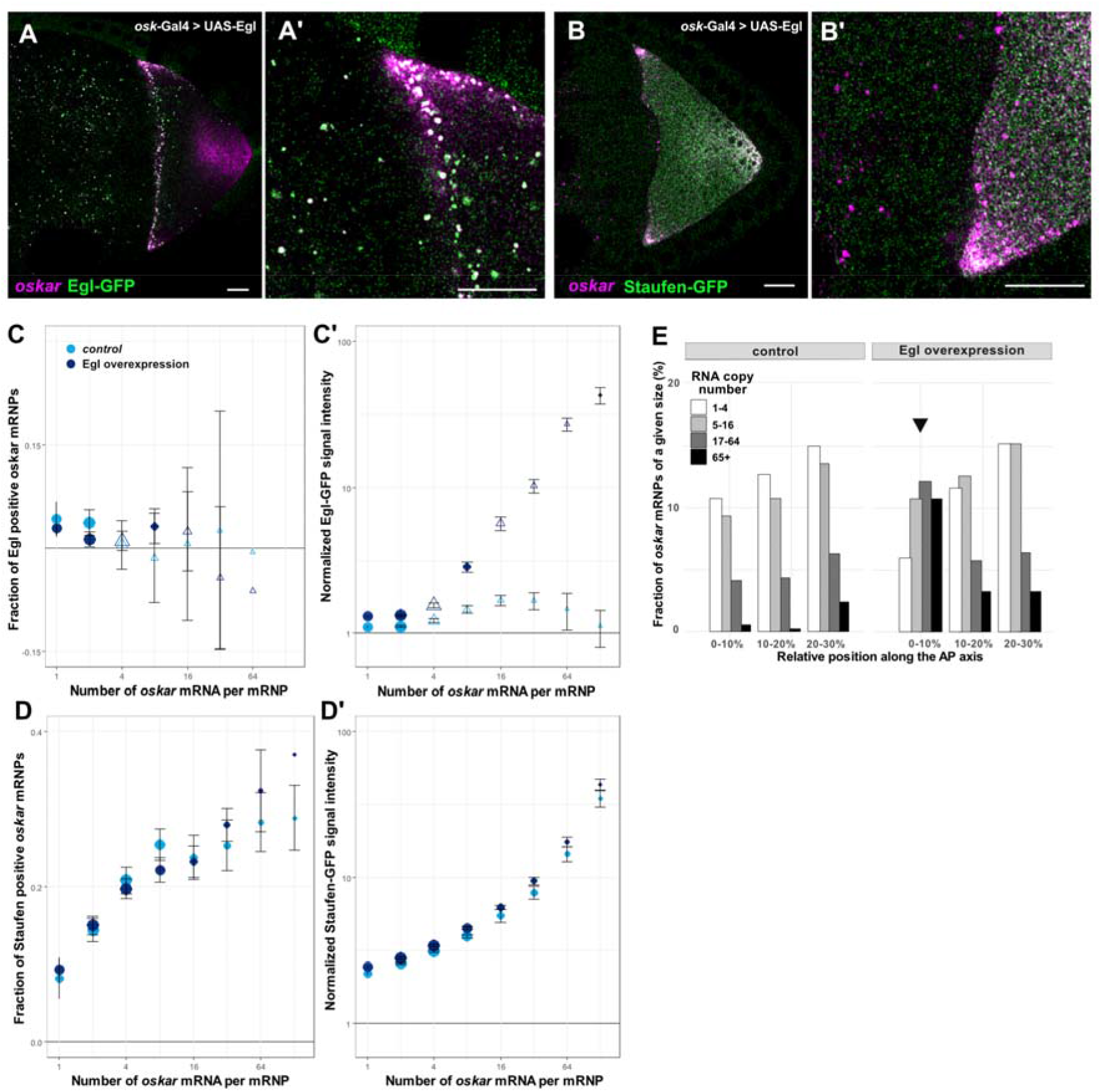
Effect of Egl overexpression on *oskar* RNP localization and composition. (A-B’) Images of *oskar* mRNA and Egl-GFP (A and A’) or Staufen-GFP (B and B’) in stage 9 egg-chambers strongly overexpressing Egl in the germline (*oskar-Gal4>UAS-Egl)*. In both genotypes, abnormally large RNPs containing *oskar* mRNA are observed in the nurse cells and in the anterior region of the oocyte. These RNPs frequently colocalize with Egl-GFP but rarely co-localize with Staufen-GFP. Scale bar represents 10 μm. (C-D’) Quantification of Egl-GFP and Staufen-GFP association with *oskar* RNPs as a function of RNA copy number. Error bars represent 95% confidence intervals and the size of the circles is proportional to the relative abundance of each category of *oskar* RNP within the overall population. Triangles indicate that the fraction of GFP-positive *oskar* RNPs is not-significantly different from zero (p>0.01, C-D’). (E) Relative distribution of *oskar* RNPs grouped by RNA content along the first 30% of the anteroposterior axis of stage 9 oocytes. In the wild-type control, less than 1% of large (65+ copies) *oskar* RNPs are close to the oocyte anterior (first bin). When Egl is overexpressed, approximately 10% of large *oskar* RNPs are present close to the oocyte anterior (first bin, arrow), while the rest of the oocyte has a distribution of *oskar* RNPs similar to the control (Fig S5A).

Taken together, these data reveal that a shared feature of *oskar* mislocalization to the anterior in the absence of Staufen, upon Egl overexpression, and in gain-of-function *BicD* mutants is the failure to release Egl from the RNPs during mid-oogenesis. Our data indicate that this release is mediated by Staufen, which interferes with the association of Egl with *oskar* RNPs, especially those with high *oskar* mRNA content.

### Dissociation of Egl does not prevent association of dynein with *oskar* RNPs

As described above, Egl functions as an adaptor molecule linking mRNA localization signals to dynein-dynactin (Navarro *et al.*, 2004; Dienstbier *et al.*, 2009; Amrute-Nayak and Bullock, 2012). However, it has been proposed that Egl is not the only means to link the motor complex to RNPs. In *ex vivo* experiments, both the *K10* and *hairy* mRNAs recruit some dynein-dynactin complexes through Egl-BicD, and others through an Egl-BicD independent mechanism, which presumably involved other RBP linkers or direct interactions between the RNA and the motor complex (Bullock *et al.*, 2006; Amrute-Nayak and Bullock, 2012; Dix *et al.*, 2013; Soundararajan and Bullock, 2014). Only those dynein-dynactin complexes interacting with Egl-BicD were capable of long-distance transport, explaining how these two adaptors promote minus-end-directed trafficking of mRNAs. How Egl influences dynein’s association with RNPs has not, however, been examined *in vivo*.

To address this question, we quantified the colocalization of *oskar* mRNA (detected by smFISH) with fluorescently tagged versions of p50/Dynamitin, a dynactin subunit, BicD and Dynein heavy chain (Dhc) in the nurse cells and oocyte. We saw little difference in association of each protein with *oskar* mRNA between the stage 9 nurse cells and oocyte (Fig. 6A-D). This contrasts with the situation for Egl, which showed a relative reduction in association with *oskar* in the stage 9 oocyte (Fig. 4D and D’). The fraction of *oskar* RNPs associated with BicD and the dynein-dynactin subunits also did not change as a function of RNA content in the oocyte (Fig 6D). Notably, the relative amount of these proteins associated with *oskar* RNPs in the oocyte was approximately double that in the nurse cells (Fig 6E). The amount of BicD and p50 associated with the RNPs in the oocyte was independent of *oskar* copy number, whereas Dhc association increased as a function of *oskar* mRNA content (Fig 6E). This result indicates that Dhc and BicD/p50 are not recruited to *oskar* RNPs by identical mechanisms.

**Figure 6:**
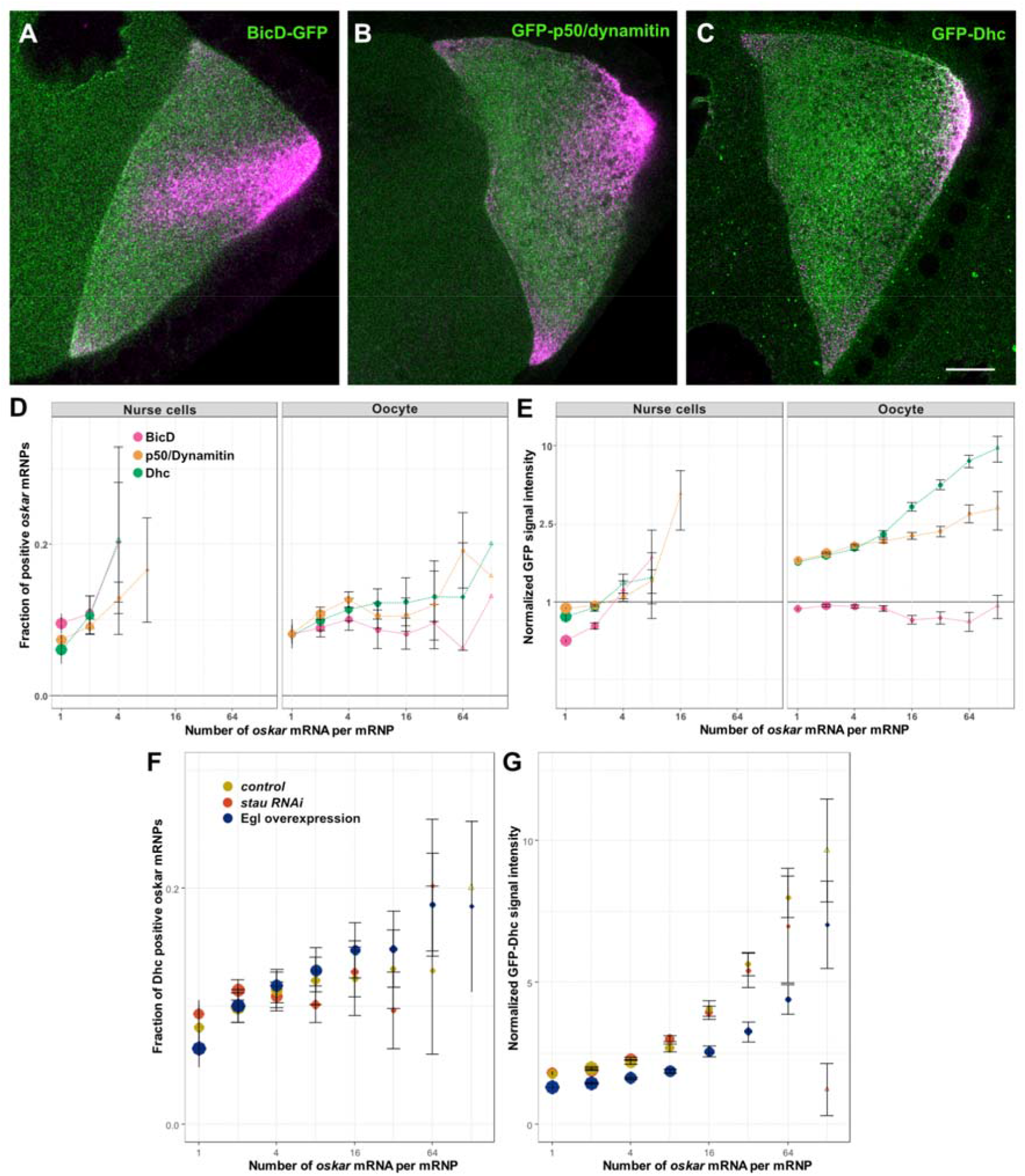
Recruitment of the dynein machinery during the transport of *oskar* RNPs. (A-C) Localization of GFP tagged versions (green) of BicD (A), p50/dynamitin (B) and Dhc (C) with respect to *oskar* mRNA (magenta) in egg-chambers during mid-oogenesis. Scale bar represents 10 μm. (D, E) Quantification of *oskar* RNP association with the indicated proteins in the nurse cells and the oocyte as a function of RNA copy number. (F, G) Quantification of *oskar* RNP association with GFP-Dhc in the oocytes of indicated genotypes. Error bars represent 95% confidence intervals and the size of the circles is proportional to the relative abundance of each category of *oskar* RNP within the overall population. Triangles indicate that the fraction of GFP-positive *oskar* RNPs is not-significantly different from zero (p>0.01, D-G).

Increased levels of Egl on *oskar* RNPs in the absence of Staufen (Fig 4E, E’ and S5E, E’) or in the presence of excess Egl (Fig 5D, D’) could in principle result in increased recruitment of the dynein motor. However, when we analyzed GFP-Dhc association with *oskar* RNPs, we found no major difference between *stau* RNAi, Egl-overexpressing, and control oocytes (Fig 6F and F’). These data indicate that Egl is not obligatory for recruitment of dynein-dynactin to a native cargo *in vivo*. Intriguingly, *stau* RNAi and Egl overexpressing oocytes also did not display an increase in the association of BicD with *oskar* RNPs (Fig 6G and G’). Thus, there appears to be an Egl-independent mechanism for recruiting BicD to these structures.

The loss of Egl from larger *oskar* RNPs during stage 9 suggests that dynein – although present – is not fully active. This notion is consistent with our earlier observation in control ooplasmic extracts that RNPs with a greater RNA content, and which therefore have less Egl bound (Fig 1B and 4D and D’), engaged in minus-end-directed motion less frequently than those with a lower RNA content: there was an approximately 4-fold plus-end dominance (77.9% vs 22.1% of plus vs minus-end directed runs) for *oskar* RNPs with more than two relative RNA units, versus an approximately 2-fold plus-end dominance (~65% vs ~35%) for smaller RNPs (Fig 1B). In summary, in stage 9 oocytes, *oskar* RNPs lacking Egl maintain their association with the dynein machinery but are defective in minus-end-directed motion.

### Staufen activity is controlled by Egl-mediated delivery of its mRNA into the oocyte

As shown above, proper *oskar* localization to the posterior pole depends on availability of Staufen protein in stage 9 oocytes. However, it is unclear how this spatio-temporal restriction of Staufen is brought about. We noticed that only trace amounts of endogenous Staufen (Fig 7C and D) or Staufen-GFP (Fig S7C) accumulated in the early arrested oocyte primordium in *egl* RNAi egg-chambers. This observation raised the possibility that Egl mediates the enrichment of Staufen in the ooplasm of wild-type egg-chambers. The normal ooplasmic accumulation of Staufen could in principle result from its transport in association with target mRNAs from the nurse cells into the oocyte. However, this seems unlikely, as we detected no association of Staufen-GFP with its main target, *oskar* mRNA, in the nurse cells (Fig 4A-B’), and both Staufen and Staufen-GFP signal accumulated in the oocyte even in the absence of *oskar* mRNA (Fig S7A-C). Alternatively, *stau* mRNA could be transported by Egl-BicD-dynein-dynactin into the oocyte, leading to accumulation of Staufen protein at this site. Supporting this notion, whereas *stau* mRNA and protein were strongly enriched in oocytes of early-stage egg chambers expressing control RNAi, we observed little or no such enrichment in egg-chambers in which *egl* was knocked down by RNAi (Fig 7B and D). Furthermore, while *stau* mRNA did not display any specific localization pattern in the ooplasm of wild-type oocytes at mid-oogenesis (Fig 7E), upon Egl overexpression the transcript was enriched at the anterior pole of the oocyte (Fig 7E’). This observation suggests that, similar to other oocyte-enriched transcripts, such as *oskar*, *gurken* and *bicoid* (Ephrussi *et al.*, 1991; Berleth *et al.*, 1988; Neuman-Silberberg and Schüpbach, 1993; Theurkauf *et al.*, 1993; Clark *et al*. 2007), *stau* mRNA is a target of Egl-dependent transport from the nurse cells by dynein.

**Figure 7:**
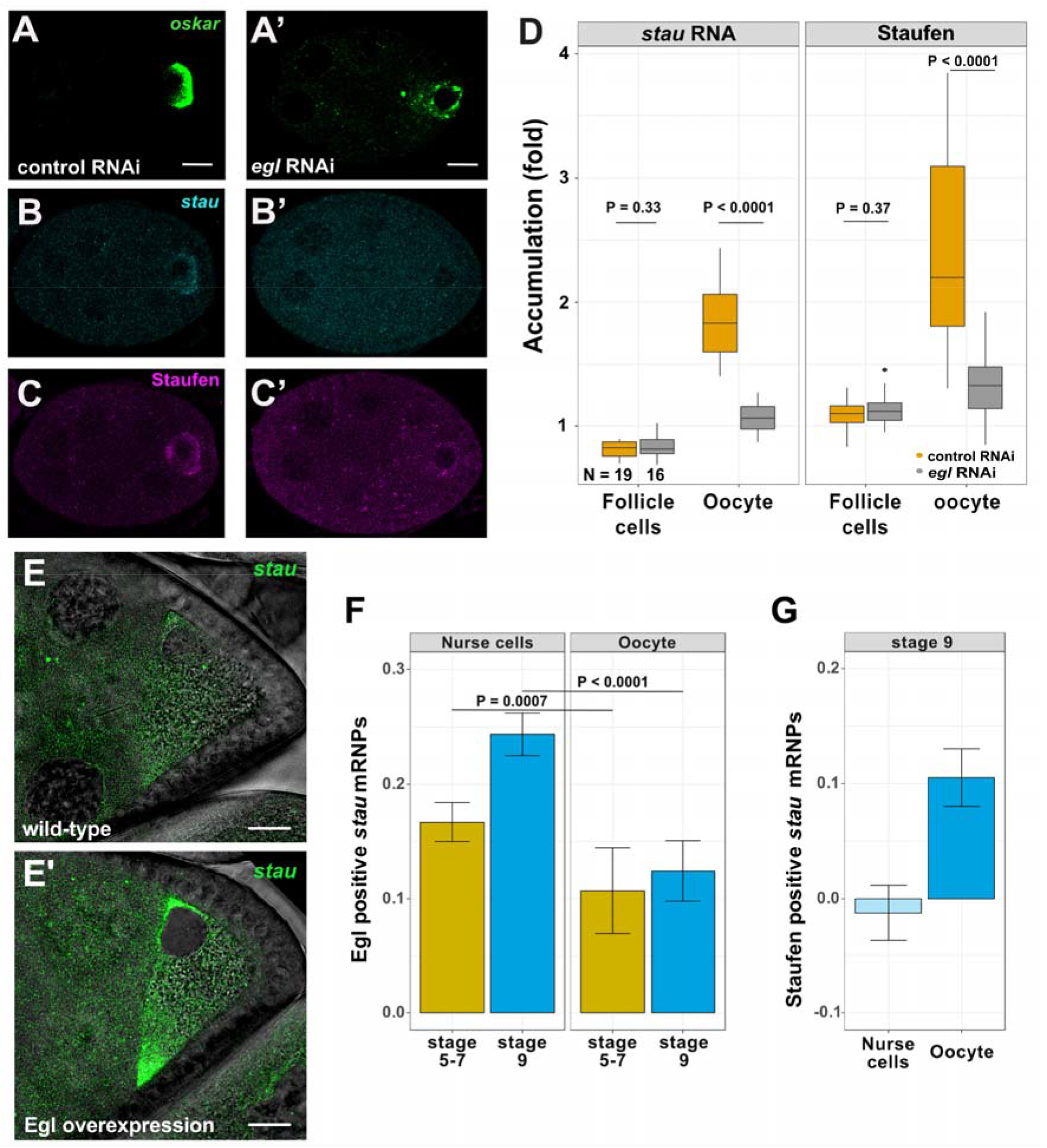
Egl promotes ooplasmic enrichment of Staufen mRNA and protein. (A-C’) Distribution of endogenous *oskar* mRNA (A, A’; green), *stau* mRNA (B, B’; cyan) and Staufen protein (C, C’; magenta) in early egg-chambers expressing control RNAi (A, B, C) or *egl* RNAi driven by *osk*-Gal4 (A’, B’, C’). Oocytes of *egl* RNAi egg-chambers contain trace amounts of *oskar* mRNA, likely due to the action of residual Egl protein. (D) Quantification of oocyte enrichment of *stau* mRNA or Staufen protein relative to the sibling nurse cells. Oocytes were identified through their enrichment of *oskar mRNA.* Enrichment of *stau* RNA or Staufen protein in the somatic follicle cells, which do not express the shRNA, is used as a control. 19 control RNAi and 16 *egl* RNAi samples were analyzed, respectively (also indicated on the panel). P-values of unpaired student’s t-test are shown. (E, E’) Localization of *stau* mRNA in wild-type and Egl-overexpressing stage 8 oocytes. (F, G) Fraction of *stau* RNPs associating with Egl in the nurse cells and in the oocyte (F), and Staufen in the oocyte (G). Scale bars represent 10 μm.

Providing further support for this notion, colocalization analysis revealed that *stau* mRNA and Egl-GFP were associated in the germline, and that this association was significantly lower in the oocyte than in the nurse cells (Fig 7F). We also detected Staufen protein in association with *stau* RNPs in stage 9 oocytes (Fig 7G), as we previously observed for *oskar* RNPs (Fig 4).

Additional experiments provided evidence that the restriction of Staufen activity to the oocyte is critical for proper RNA localization and oogenesis. In contrast to the direct expression of GFP-Staufen from the maternal tubulin promoter, which results in nurse cell accumulation of the protein only from mid-oogenesis onwards (Fig S7I), strong overexpression of GFP-Staufen with highly active maternal Gal4 drivers results in high levels and aggregation of the protein in the nurse cells of early egg-chambers (Fig S7E-H). Ectopic GFP-Staufen expression in this manner resulted in failure of *oskar*, as well as *bicoid*, mRNA enrichment in the oocyte (Fig S7E-F’’). These egg-chambers also displayed oocyte polarity defects and fragmentation of the karyosome (Fig S7G and H), which are hallmarks of defects in localization of mRNA cargoes for dynein (Neuman-Silberberg and Schüpbach, 1993; Jenny *et al.*, 2006).

Taken together, these data indicate that the Egl-dependent transport of *stau* mRNA into the oocyte by dynein underlies ooplasmic enrichment of Staufen protein, which in turn inactivates the machinery for minus end-directed transport of *oskar* RNPs in the stage 9 oocytes. This constitutes a feed-forward mode of regulation, whereby the dynein transporter system ensures the localized production of its inhibitor, leading to its own inactivation. This predefined program thereby allows kinesin-1-dependent delivery of *oskar* mRNA to the oocyte posterior.

## DISCUSSION

The activity of motor molecules is essential for establishing proper localization of mRNA molecules, which in turn underlies spatiotemporal restriction of protein function (St Johnston, 2005; Martin and Ephrussi, 2009; Mofatteh and Bullock, 2017; Abouward and Schiavo, 2021). The dynein and kinesin microtubule motors play key roles in positioning of mRNAs in many systems, including by acting sequentially on the same RNP species (Baumann *et al.*, 2012; Gagnon *et al.*, 2013; Mofatteh and Bullock, 2017; Turner-Bridger *et al.*, 2018). However, it is unclear how the opposing activities of these motors are coordinated during RNP trafficking.

Here we use the tractable *Drosophila* egg-chamber to reveal a mechanism for spatiotemporal control of dynein and kinesin-1 activity. Central to this system are two dsRBPs, Staufen and Egl. Several genetic interaction studies have shown previously that these proteins have opposing activities in the context of *oskar* mRNA localization in the oocyte and anteroposterior patterning (Mohler and Wieschaus, 1986; Navarro *et al.*, 2004) and both proteins were shown to form a complex with *oskar* (Laver *et al.*, 2013; Sanghavi *et al.*, 2016). Recent work by Mohr and colleagues (Mohr *et al.*, 2021), which was published while this manuscript was in preparation, further corroborated the ability of Egl to antagonize localization of *oskar* to the posterior of the oocyte. Mohr *et al.* also investigated the potential interplay of Staufen and Egl *in vitro* with RNA binding assays. It was shown that *in vitro* Egl binds to a truncated version of stem-loop 2 in the *oskar* 3’ UTR (SL2, (Jambor *et al.*, 2014) that they termed the Transport and Anchoring Signal (TAS) (Mohr *et al.*, 2021). The TAS partially overlaps with one of the Staufen Recognized Structures (Laver *et al.*, 2013) which are important for Staufen binding *in vitro* and for proper *oskar* mRNA localization *in vivo* (Mohr *et al.*, 2021). These data suggest that Staufen could antagonize Egl function by interfering with binding of the latter protein to *oskar* mRNA. However, Mohr et al. did not directly test this possibility. We found that knocking down Staufen increases the association of Egl with the mRNA, demonstrating a role for Staufen in antagonizing association of Egl with *oskar* RNA. The results of our *in vitro* motility assays are also consistent with this scenario, as RNA binding to Egl is a prerequisite for full dynein activity within this minimal RNP (McClintock *et al.*, 2018; Sladewski *et al.*, 2018). It is conceivable that the transport and anchoring functions ascribed to TAS are two facets of the same underlying molecular mechanism, as increased dynein activity in the absence of Staufen would drive enrichment of *oskar* RNPs at microtubule minus ends that are nucleated at the anterior cortex (Januschke *et al.*, 2006).

Our study additionally reveals through smFISH-based colocalization analysis in egg-chambers how the interactions of Staufen and Egl with *oskar* mRNA are orchestrated in time and space. When Staufen levels are low, such as in early oogenesis, amounts of Egl per RNP scale with *oskar* RNA content. During stage 9, this scaling is lost in the oocyte and the relative amount of Egl on *oskar* RNPs decreases as Staufen is recruited to *oskar*. Our data indicate that Staufen-mediated displacement of Egl is a critical step in switching to kinesin-1-based trafficking of *oskar* mRNA to the posterior pole. This switch is also likely to involve the activities of exon junction complex components, which are required for posterior localization of *oskar* (Newmark and Boswell, 1994; Hachet and Ephrussi, 2001, 2004; Mohr, Dillon and Boswell, 2001; van Eeden *et al.*, 2001; Palacios *et al.*, 2004; Zimyanin *et al.*, 2008; Ghosh *et al.*, 2012). Our study provides a framework for understanding how the activities of these factors are coordinated with that of Staufen.

Intriguingly, whilst an excess of Staufen can dissociate Egl from *oskar* RNPs, increasing Egl concentration does not overtly affect Staufen recruitment with these structures. This observation might be due to the five-to-one excess of SRSs over the TAS in each *oskar* molecule (Mohr *et al.*, 2021), which could mask the loss of Staufen binding to the TAS-proximal SRS.

We also found that dissociation of Egl from *oskar* RNPs in the stage 9 oocyte does not detectably alter the amount of dynein on these structures. This work lends *in vivo* support to the notion that additional proteins or RNA sequences can recruit dynein and dynactin in an inactive state to RNPs (Bullock *et al.*, 2006; Amrute-Nayak and Bullock, 2012; Dix *et al.*, 2013; Soundararajan and Bullock, 2014). Presumably, Egl recruits only a small fraction of the total number of dynein complexes on *oskar* RNPs or activates the motility of complexes that are also linked to the RNA by other factors. Unexpectedly, we reveal that the bulk association of BicD with *oskar* RNPs is also not dependent on Egl, pointing to an additional mechanism for recruiting BicD, presumably in the autoinhibited state, that prevents dynein activation (Hoogenraad *et al.*, 2003; Liu *et al.*, 2013). While the relative amount of the dynein machinery associated with *oskar* RNPs is greater in the oocyte than the nurse cells, dynein’s association with *oskar* RNPs in the oocyte does not scale with RNA content. How the amount of dynein recruited to these RNPs could be limited is unclear. However, this mechanism could be a means to prevent sequestration of this multi-functional motor to a very abundant cargo (~0.5-1 million copies of *oskar* mRNA per oocyte (Little *et al.*, 2015)).

Whilst our data build a strong case that a key function of Staufen in *oskar* mRNA localization is to limit dynein activity by displacing Egl, several lines of evidence suggest that this is not its only role in this context. We found that Staufen had a partial inhibitory effect on minus-end-directed motility of purified dynein-dynactin complexes activated in an Egl-independent manner by a constitutively active truncation of a BicD protein. Understanding the mechanistic basis of this effect will be the goal of future studies. Moreover, Mohr *et al.* (Mohr *et al.*, 2021) found that the SRSs that are not proximal to the TAS in the sequence of *oskar* also contribute to posterior localization of *oskar* mRNA in a Staufen-dependent manner. Although it is possible that these elements are close enough to the binding site for Egl-BicD-dynein-dynactin in the folded RNA molecule to interfere directly with the assembly or activity of the complex, they could also regulate *oskar* mRNA distribution through an independent mechanism. Other observations in our study hint at other roles of Staufen. We observed that when Staufen is depleted motile *oskar* RNPs tend to have a lower RNA content, suggesting an additional function of the protein in RNA oligomerization. Furthermore, whilst the magnitude of the effect was much smaller than for minus-end-directed motion, there was increased plus-end-directed velocity of a subset of *oskar* RNPs in *ex vivo* motility assays when Staufen was disrupted. Although this observation could reflect dynein’s ability to promote kinesin-1 activity (Hancock, 2014), it is also possible that Staufen directly tunes the activity of the plus end-directed motor.

An important question raised by our analysis of Staufen's effects on *oskar* mRNA transport is how the timing and location of this process is controlled. We provide evidence that this key aspect of *oskar* mRNA localization is based on another mRNA localization process in which Egl, as part of s *tau* RNPs, is responsible for the enrichment of *stau* mRNA in the developing oocyte. We propose this mechanism constitutes a feed-forward type of switch, whereby the activity of the dynein-mediated transport machinery deploys its own negative regulator to a distant location. The resultant increase in levels of ooplasmic Staufen levels in the ooplasm prevents dynein-mediated transport of *oskar* to the anterior of the oocyte, while the almost complete absence of the protein from the nurse cells is likely to be important for uninterrupted transport of *oskar* RNPs into the oocyte by dynein during early- and mid-oogenesis. Presumably, *stau* translation is suppressed during transit into the oocyte or the protein is translated en route but only builds up to meaningful levels where the RNA is concentrated in the oocyte. Staufen protein might also modulate its own localization in the ooplasm by antagonizing the association of *stau* mRNA with Egl, and thereby its minus end-directed motility. Consistent with this notion, the level of Egl on *stau* RNPs declines at a similar stage to when Staufen inhibits the association of Egl with *oskar* RNPs. As Staufen controls localization and translation of other mRNAs in the maturing oocyte (St Johnston *et al.*, 1991; Ferrandon *et al.*, 1994), mRNA-based regulation of the protein distribution in the egg-chamber is likely to have functions that extend beyond orchestrating *oskar* mRNA localization. Given the functional conservation of Staufen protein (Heber *et al.*, 2019) and the observation that the mRNA encoding mStau2 is localized in dendrites of mammalian neurons (Zappulo *et al.*, 2017), it is plausible that the feed-forward loop established by *stau* RNA localization during *Drosophila* oogenesis is an evolutionarily conserved process that controls RNA trafficking and protein expression in polarized cells.

## METHODS

### Fly strains

To knock down *stau* and *egl* RNA levels, we used the P{TRiP.GL01531} (FBal0281234) and the P{TRiP.HM05180} (FBal0240309) transgenic lines. A TRiP line against the *w* gene (P{TRiP.GL00094} - FBal0262579) was used as a negative control. The following mutant lines were used to disrupt or modify gene function: *stau*[D3] (FBal0016165) and *stau*[R9] (FBal0032815) alleles in heterozygous combination; *egl*[1] (FBal0003574) and *egl*[2] (FBal0003575); *BicD*[1] (FBal0001140) and *BicD*[2] (FBal0001141); *osk*[A87] (FBal0141009) in combination with *Df(3R)p-XT103* (FBab0002842) to remove *oskar* mRNA from egg-chambers. *w*[1118] (FBal0018186) was used as the wild-type control.

To overexpress Staufen, we used the αTub67C:GFPm6-Staufen (FBal0091177), αTub67C:RFP-Staufen (Zimyanin *et al.*, 2008) transgenic lines and a P{UASp-Staufen} transgene inserted onto the 3^rd^ chromosome (kind gift of F. Besse).

To overexpress unlabelled Egl protein, we used the P{UASp-egl.B} transgene (FBtp0022425) inserted on the X or the 3^rd^ chromosome. To label proteins of interest, we used the following fluorescently tagged reporter lines: P{Staufen-GFP} inserted on the 3^rd^ chromosome, expressing Staufen-GFP under control of the endogenous *stau* promoter (kind gift of F. Besse); P{tub-egl.EGFP} (FBtp0041299) inserted on the 3^rd^ chromosome, driving expression of Egl-GFP; P{UASp-BicD.eGFP} (FBtp0069955) and P{UAS-DCTN2-p50::GFP} (FBtp0016940) transgenic insertions to label BicD and Dynamitin, respectively. To drive the expression of the TRiP RNAi lines and other UASp constructs in the female germline, one copy of P{osk-Gal4} (FBtp0083699) inserted onto either the 2^nd^ or the 3^rd^ chromosome was used. For moderate overexpression of UASp-GFP-Staufen, we used one copy of P{matα4-GAL-VP16} (FBtp0009293) inserted onto the 2^nd^ chromosome.

The endogenous Dhc locus (*Dhc64C*) was tagged with the EmeraldGFP coding sequence to generate a GFP-Dhc expressing fly line according to protocols of the flyCRISPR website (Gratz *et al.*, 2013; O’Connor-Giles, Wildonger and Harrison, 2014) (https://flycrispr.org/). The locus was targeted using CRISPR/Cas9 and a guide RNA targeting the following sequence: 5’ GAGTCACCCATGTCCCACAA. The introduced double-stranded break was repaired by homologous recombination using an in-frame EmeraldGFP coding sequence flanked by two ~ 700 bp long homology arms targeting around the Dhc translational initiation codon. F1 generation embryos were screened for GFP fluorescence to identify individuals with a modified genome. Flies homozygous for GFP-Dhc are viable and fertile.

All stocks were raised on normal cornmeal agar at 25C▫C, and females were fed with wet yeast overnight before harvesting their ovaries.

### *Ex vivo* motility assay of *oskar* RNPs

The *ex vivo* motility analysis of *oskar* RNPs were carried out as described in (Gáspár *et al.*, 2017). Briefly, control and *stau* RNAi ovaries expressing mCherry-alpha-tubulin and *oskMS2*-GFP were dissected and transferred onto silanized coverslips in a drop of BRB80. Several stage 9 egg-chambers were isolated and pulled under Voltalef 10S halocarbon oil on the same coverslip. Under the oil, the nurse cells were removed by laceration using two fine tungsten needles. The isolated oocyte still in a “sack” of follicle cells were spatially separated from the remnants of nurse cell cytoplasm and with a gentle prick on the oocyte anterior the ooplasm was released onto the coverslip by a continuous slow pulling on the posterior of the oocyte-follicle “sack”. Once several such preps were created, the ooplasmic extracts were imaged using a Leica 7000 TIRF microscope with 100x oil (NA=1.4) objective to visualize *oskMS2*-GFP and mCherry labeled microtubules. Time-lapse series were collected and analyzed as described in (Gáspár *et al.*, 2017).

To analyze the relative mRNA content of *oskMS2*-GFP RNPs, a series of Gaussian functions was fitted to the GFP signal intensity distribution in each time-lapse series individually using the mixtools package in R (Benaglia *et al.*, 2009; R Core Team, 2014). The smallest *μ* value of Gaussian fits was used to represent a single unit of RNA and each RNP was normalized to this value.

### RNA co-immunoprecipitation (RIP) from ovarian lysate

RIP was carried out as described in (Gáspár *et al.*, 2017). Briefly, ovaries from 50 flies were dissected in BRB80 (80 mM PIPES, 1 mM MgCl_2_, 1 mM EGTA, pH 6.8) and lysed in Pre-XL buffer (20 mM Tris-Cl, 150 mM KCl, 4 mM MgCl_2_, 2xPIC, 1 mM PMSF, pH 7.5; supplemented with 40U of RiboLock RNase Inhibitor per 100μL lysate). Ovaries were ground using a pestle and centrifuged for 1 min at 500 × g. The supernatant was extracted and crosslinked at 0.3J/cm^2^. The lysate was equalized with 1 volume of Pre-XL buffer, 1 volume of RIPA (10 mM Tris/Cl pH 7.5; 150 mM NaCl; 0,5 mM EDTA; 0,1% SDS; 1% Triton X-100; 1% Deoxycholate, 0.09% Na-Azide) buffer and 8 volumes of low salt buffer (20 mM Tris-Cl, 150 mM KCl, 0.5 mM EDTA, 1 mM PMSF). GFP-Trap®_MA beads were washed with low-salt buffer and blocked for 60 mins at room temperature in Casein Blocking Buffer (Sigma-Aldrich) supplemented with 50 μg/mL Heparin. Lysate was incubated with beads for 90 minutes at 4°C. The beads were then washed six times with high salt buffer (20 mM Tris-Cl, 500 mM NaCl, 1 mM EDTA, 0.1% SDS, 0.5 mM DTT, 1xPIC, 1mM PMSF) and twice with PBT (PBS + 0.1% Triton). Endogenous RNA cross-linked to bait protein was recovered from the beads using the QuickRNA Microprep Kit (Zymo Research). Complementary DNA was synthesized using the SuperScript III First-Strand Synthesis kit (Invitrogen) and used as template for PCR using *oskar* and *bicoid* primers.

### Single molecule fluorescent hybridization (smFISH)

Single molecule FISH was carried out as described previously (Gáspár *et al.*, 2017; Gaspar, Wippich and Ephrussi, 2017; Heber *et al.*, 2019). Briefly, ssDNA oligonucleotides complementary to *oskar* and *stau* mRNAs (Table S1) were mixed and labeled with Atto532, Atto565 or Atto633 according to the protocol described in Gaspar et al., 2017 (Gaspar, Wippich and Ephrussi, 2017).

*Drosophila* ovaries expressing fluorescent reporter transgenes were dissected in 2% PFA, 0.05% Triton X-100 in PBS and fixed for 20▫min. The fixative was removed by two 5 minute washes in PBT (PBS▫+▫0.1% Triton X-100, pH 7.4). Ovaries were pre-hybridized in 200▫μL 2▫×▫HYBEC (300▫mM NaCl, 30▫mM sodium citrate pH 7.0, 15% ethylene carbonate, 1▫mM EDTA, 50▫μg per mL heparin, 100▫μg per mL salmon sperm DNA, 1% Triton X-100) for 10▫min at 42▫°C. Meanwhile, the 50 μL probe mixture (5CnM per individual oligonucleotide) was prepared and pre-warmed to hybridization temperature. After 10 minutes of pre-hybridization, the probe mix was mixed into the pre-hybridization mixture. After 2▫hours of hybridization at 42▫°C, the unbound probe molecules were washed out of the specimen by two washes in pre-warmed HYBEC and a final wash in PBT at room temperature. To boost signal intensity for the protein enrichment analysis of Staufen-GFP in early egg-chambers, we incubated the ovaries in GFP-Booster (Chromotek) diluted 1:2000 in PBT for 1 hour, then washed the sample three times for 10 minutes in PBT. Ovaries were mounted in Vectashield and processed for smFISH analysis.

### Microscopy

Imaging of oocytes was carried out on a Leica TCS SP8 confocal laser scanning microscope using a 20x dry (NA▫=▫0.75) objective for imaging the RNA distribution in the oocytes and a 63x oil immersion (NA▫=▫1.4) objective for analysis of RNA or Staufen-GFP protein enrichment in early oocytes and for colocalization analysis between RNPs and the fluorescently labeled Staufen and Egl proteins.

Imaging of reconstituted RNPs by TIRF microscopy was performed as described previously (McClintock *et al.*, 2018) using a customized Nikon TIRF system controlled by Micro-manager (Edelstein *et al.*, 2014), with the exception of data presented in Figures 2H and I, and Figure S3. For these experiments, movies were acquired with 100-ms exposures at 4 frames/s for a total of 3 min on a Nikon Eclipse Ti2 inverted TIRF system equipped with a Nikon 100x Apo TIRF oil immersion objective (NA = 1.49) and Photometrics Prime 95B CMOS camera. TMR-dynein was imaged using the 561 nm laser line with a Chroma ET-561 nm Laser Bandpass Set (ET576lp and ET600/50m) and ET605/52m emission filter.

### RNA distribution analysis

Analysis of *oskar* and *bicoid* mRNA distribution was carried out as described in (Gaspar *et al.*, 2014; Gáspár *et al.*, 2017; Heber *et al.*, 2019). Briefly, we manually defined the outlines of the oocytes and their anteroposterior (AP) axis, using a custom-made ImageJ plug-in, the smFISH signal was redistributed into a 100▫×▫100 matrix. Each column of the matrix represents the relative amount of signal found under 1% of the AP axis length, with anterior on the left (column 1) and posterior on the right (column 100). The matrices from different oocytes of the same genotype and stage were averaged to obtain an average view RNA localization. The center-of-mass (relative to the geometric center of the oocyte) was determined and compared statistically using a Kruskal–Wallis test followed by pairwise Mann–Whitney U test against the control.

### RNA and protein enrichment analysis

The boundaries of the somatic follicle cells, nurse cells and oocyte were manually defined, and the fluorescence intensity of smFISH or GFP signal in these compartments was measured by ImageJ. The extent of enrichment of RNA and protein in the somatic follicle cells and the oocyte was obtained by normalizing the measured fluorescence intensity values to the corresponding values obtained for the nurse cells.

### Colocalization analysis between RNPs and fluorescent reporters

Analysis was carried out as described in (Gáspár *et al.*, 2017; Heber *et al.*, 2019). Briefly, images for colocalization analysis were deconvolved using Huygens Essential (https://svi.nl) and segmented using a custom ImageJ (Schneider, Rasband and Eliceiri, 2012) plug-in. Nearest-neighbor pairs between RNPs and fluorescent reporters were identified, their position and signal intensities were extracted from the nurse cell and oocyte compartments, excluding any nuclear areas in the field of view. Quantification of mRNA copy number per RNP and normalization of fluorescent reporter signal was carried out by fitting multiple Gaussian functions to the corresponding signal intensity distributions taken from the nurse cells using the mixtools package in R (Benaglia *et al.*, 2009; R Core Team, 2014). The *μ* value of Gaussian fit that described the largest portion of the distribution in the nurse cells (almost the lowest value of all fitted μ values) was taken as the signal intensity of one unit (for RNPs the intensity of a signal mRNA molecule). These unit values were used to normalize raw signal intensities. RNPs were clustered based on this normalized intensity under the following rule: [2^i^:2^i+1^), i ∈ [0:8], i.e., 1, 2:3, 4:7, 8:15, etc. The observed nearest-neighbor colocalization frequencies were computed for each cluster and were compared to the expected colocalization frequencies (governed by the object-densities, determined in randomized object localizations). Similarly, the mean, normalized intensity of colocalizing fluorescent reporter molecules was calculated for each cluster. Correlation between RNA content and the normalized mean fluorescent reporter intensity was tested and compared using least-squares means analysis in R (Lenth, 2016). We typically analyzed 3000-80000 *oskar* RNPs imaged in 4-6 egg-chambers collected from 3-4 females per condition.

### Statistical analysis

Statistical analyses were performed as indicated in the figure legends using R (R Core Team, 2014), RStudio (www.rstudio.com), and GraphPad Prism version 9.1.1 for MacOS (GraphPad Software, San Diego, California USA, www.graphpad.com). All graphs were plotted by the ggplot2 library in R (Wickham, 2016) and GraphPad Prism version 9.1.1 for MacOS (GraphPad Software, San Diego, California USA, www.graphpad.com).

### Protein expression, purification, and fluorescent labeling

Recombinant dynein (human), Egl (*Drosophila*), BicD (*Drosophila*), and BicD2N (mouse) were expressed, purified, and fluorescently labeled as described previously (Hoang *et al.*, 2017; McClintock *et al.*, 2018). Briefly, complexes of the complete dynein holoenzyme,the coexpressed Egl/BicD complex, and BicD2N were expressed in *Sf*9 insect cells and purified by affinity chromatography using ZZ-tags on the Dynein heavy chain, Egl, or BicD2N, respectively. Fluorescent labeling of the Dynein heavy chain and BicD2N with SNAP-Cell TMR Star or SNAP-Surface Alexa Fluor 647 via N-terminal SNAPf tags was performed either on-column prior to elution from the IgG affinity resin by cleavage of the ZZ epitope by TEV protease (dynein) or in solution following TEV-cleavage (BicD2N). All complexes were further purified by FPLC-based gel-filtration chromatography, using either TSKgel G4000SWxl (dynein) or Superose 6 Increase 3.2/300 GL (Egl/BicD) columns. The dynactin complex (porcine) was purified natively from pig brains as described previously (Schlager *et al.*, 2014; Urnavicius *et al.*, 2015). Extracts were clarified and fractionated by FPLC using SP Sepharose (cation exchange, ~250ml bed volume), Mono Q 4.6/100 PE (anion exchange), and TSKgel G4000SW (gel-filtration) columns.

Recombinant Staufen (*Drosophila*) was cloned as fusion protein with an N-terminal His_6_-rsEGFP2-tag and a C-terminal His_6_-tag in pFastBacDual between the KpnI and HindIII sites and expressed in *Sf*21 insect cells using the Bac-to-Bac expression system (Thermo Fisher). The protein was purified by affinity chromatography on HisTrap FF in 1x PBS + 880 mM NaCl, 400 mM arginine, 10 mM - 200 mM imidazole, 2 mM DTT and HiTrap Heparin HP in 40 mM Bis-Tris pH 7.5, 150 mM NaCl – 1000 mM NaCl, 40 mM arginine, 2 mM DTT and a final gel-filtration on Superdex200 Increase in a final buffer of 40 mM Bis-Tris pH 7.5, 150 mM NaCl, 2 mM DTT.

In all cases, eluted protein was dispensed into single-use aliquots, flash-frozen in liquid nitrogen, and stored at −80°C.

### RNA synthesis and purification

The *oskar* RNA used for the reconstitution assay was uncapped and synthesized *in vitro* using the MEGAscript T7 Transcription Kit (Ambion). The RNA was transcribed from a linearized plasmid DNA template containing an ~529-nt region of the *osk* 3’UTR previously defined as region 2+3 that includes the oocyte entry signal (OES) and promotes the localization of reporter transcripts to the developing oocyte (Jambor *et al.*, 2014). For fluorescent labeling of transcripts, Cy5 UTP or ChromaTide Alexa Fluor 488 UTP was included in the transcription reaction with a 2-fold or 4-fold excess of unlabelled UTP, yielding transcripts with an average of ~5 or ~14 fluorophores per RNA molecule, respectively. Excess fluorescent UTP was removed using 2 successive rounds of Sephadex G-50 desalting spin columns per transcription reaction. Transcripts were subsequently purified by ammonium acetate/ethanol precipitation and resuspension in nuclease-free water. RNA was dispensed into single-use aliquots and stored at −80°C.

### *In vitro* RNP reconstitution assays

TIRF-based single-molecule reconstitution assays were performed as described previously (McClintock *et al.*, 2018). Briefly, taxol-stabilized porcine microtubules were polymerized with a mixture of unlabelled tubulin, HiLyte 488 tubulin and biotin-conjugated tubulin, and immobilized in imaging flow chambers by streptavidin-based linkages to biotin-PEG passivated cover slips. Assembly mixes containing relevant combinations of dynein, dynactin, Egl, BicD, BicD2N, *oskar* RNA, and Staufen were then diluted to concentrations suitable for imaging of single molecules (~2.5 nM for dynein), applied to the flow chamber and subsequently imaged by TIRF microscopy. For assays testing the effect of Staufen on RNA or dynein motility, the complexes were assembled for 45-60 minutes on ice with components diluted in GF150 buffer (25 mM HEPES pH 7.3, 150 mM KCl, 1 mM MgCl_2_, 5 mM DTT, 0.1 mM MgATP, 10% glycerol) to concentrations of 100 nM TMR-dynein, 200 nM dynactin, 500 nM Egl/BicD or BicD2N, and 1 μM *oskar* RNA. This assembly mix was then diluted 1:1 with either ~4 μM rsEGFP2-Staufen or Staufen storage buffer (40 mM Bis-Tris pH 7.5, 150 mM NaCl, 2 mM DTT) and incubated for at least a further 15 minutes on ice. Just prior application to the imaging chamber, this mixture was further diluted 20-fold in Motility Buffer (30 mM HEPES pH 7.3, 50 mM KCl, 5 mM MgSO_4_, 1 mM EGTA pH 7.3, 1 mM DTT, 0.5 mg ml^-1^ BSA, 1 mg ml^-1^ a-casein, 20 μM taxol) with added MgATP (2.5 mM final concentration) and 1x oxygen scavenging system (1.25 μM glucose oxidase, 140 nM catalase, 71 mM 2-mercaptoethanol, 25 mM glucose final concentrations) to reduce photobleaching. For assays testing the effect of Egl/BicD on RNA binding and motility on microtubules, the complexes were assayed as above except that primary assembly mixes consisted of 100 nM Alexa Fluor 647-dynein, 200 nM dynactin, 500 nM Egl/BicD or Egl/BicD storage buffer (GF150), and 1 μM *oskar* RNA, which was subsequently diluted 1:1 in Staufen storage buffer and processed as above. Binding and motility of RNA and dynein in the in vitro reconstitution assays were manually analyzed by kymograph in FIJI (Schindelin *et al.*, 2012) as described previously (McClintock *et al.*, 2018). Briefly, microtubule binding events were defined as those lasting for a minimum of 3 continuous frames of acquisition and processive events were defined as those that exhibited unidirectional movement of at least 5 pixels (525-550 nm depending on the microscope system) during this time, regardless of velocity.

## ACKNOWLEDGEMENTS

We thank the EMBL Advanced Light Microscopy and Gene Core Facilities for their support. We acknowledge Alessandra Reversi (EMBL *Drosophila* Injection Service) for fly transgenesis and thank Florence Besse for *Drosophila* strains and Dierk Niessing for reagents for recombinant Staufen expression. Work in S.L.B.'s group is supported by the Medical Research Council, as part of United Kingdom Research and Innovation (also known as UK Research and Innovation) [MRC file reference number MC_U105178790]. For the purpose of the MRC open access policy, the authors have applied a CC BY public copyright licence to any Author Accepted Manuscript version arising. M.A.M. is supported by a BBSRC project grant (BB/T00696X/1). L.-J.P. is supported by a FOR2333 grant of the Deutsche Forschungsgemeinschaft (Germany) to A.E. A.E. gratefully acknowledges the support of the European Molecular Biology Laboratory.

## AUTHOR CONTRIBUTIONS

I.G. and M.A.M. designed experiments. I.G., L.J.P., M.A.M. and S.H. carried out the experiments and analyzed the data. S.L.B. and A.E. supervised the work. All authors discussed the data and contributed to manuscript preparation.

## SUPPLEMENTARY FIGURES

Video S1 and S2: *Ex vivo* imaging of *oskMS2*-GFP RNPs in extracts of control (Video S1) and *stau* RNAi (Video S4) stage 9 oocytes. *oskar* RNPs are shown in green, microtubule plus tips are labeled by EB1-mCherry (magenta). Note the frequent runs of dim *oskar* RNPs in the *stau* RNAi condition (Video S2)

Video S3 and S4: Live cell imaging of *oskMS2*-GFP RNPs in control (Video S3) and *stau* RNAi (Video S4) stage 9 oocytes. Anterior (left) and posterior (right) regions of the same oocyte are shown. *oskMS2*-GFP signal is rendered in blue-yellow to allow visual appreciation of dim (blue) and bright (yellow) *oskar* RNPs.

Table S1: ssDNA oligos used to synthesize smFISH probes targeting the long 3’ UTR of *stau* A/B isoforms or the 5’ extended regions of *stau* A/C isoforms.

---

**Figure S1:**
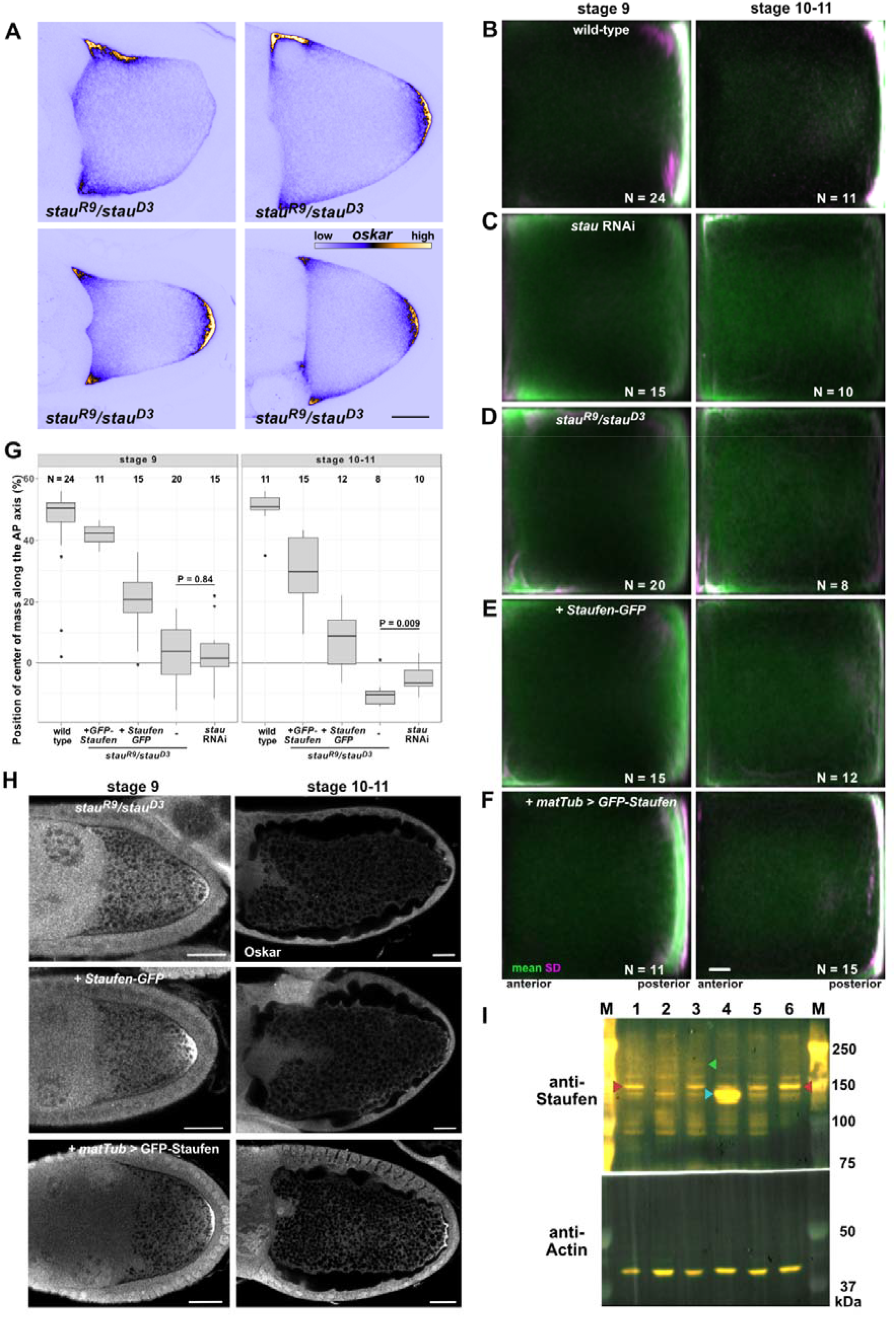
Expression and effect of the different Staufen transgenes on *oskar* mRNA localization. (A) *oskar* RNA localization (shown in blue/yellow) in stage 9 *stau^R9^/stau^D3^* mutant oocytes. Scale bars represent 20 μm. (B-F) Mean (green) and variance (SD, magenta) of *oskar* RNA distribution in stage 9 and stage 10-11 oocytes. Numbers indicate the number of oocytes analyzed for each condition and scale bar represents 10% of anteroposterior axis length. Anterior is to the left, posterior is to the right. (G) Position of *oskar* center-of-mass along the AP axis in stage 9 and stage 10-11 oocytes for each condition. 0 is the geometric center of the oocyte, with the posterior pole located at 58%. (H) Oskar protein expression in Staufen null oocytes coexpressing transgenic Staufen-GFP or GFP-Staufen. Scale bars represent 20 μm. (I) Western blot detection of Staufen in wild-type (lane 1), *stau^R9^/stau^D3^* (2), Staufen-GFP (3), GFP-Staufen (4), *stau* RNAi (5) and control RNAi (6) ovarian lysates. Note the similar distribution of *oskar* in *stau^R9^/stau^D3^* and in *stau* RNAi oocytes (C, D and G), despite residual Staufen expression in Staufen RNAi ovarian lysates (I, lanes 5 and 6). The overexpressing GFP-Staufen (I, lane 4) transgene largely rescues *oskar* mislocalization (F and G) and Oskar protein expression defects (H) observed in Staufen null mutants. The Staufen-GFP transgene, expressed at low levels (I, lane 3), rescues *oskar* RNA localization at stage 9, but the RNA is not maintained at the posterior at stages 10-11 (E and G), likely due to insufficient Oskar protein expression at the posterior (H), which is essential for *oskar* mRNA anchoring at the oocyte posterior during the later stages (Vanzo and Ephrussi, 2002).

**Figure S2.**
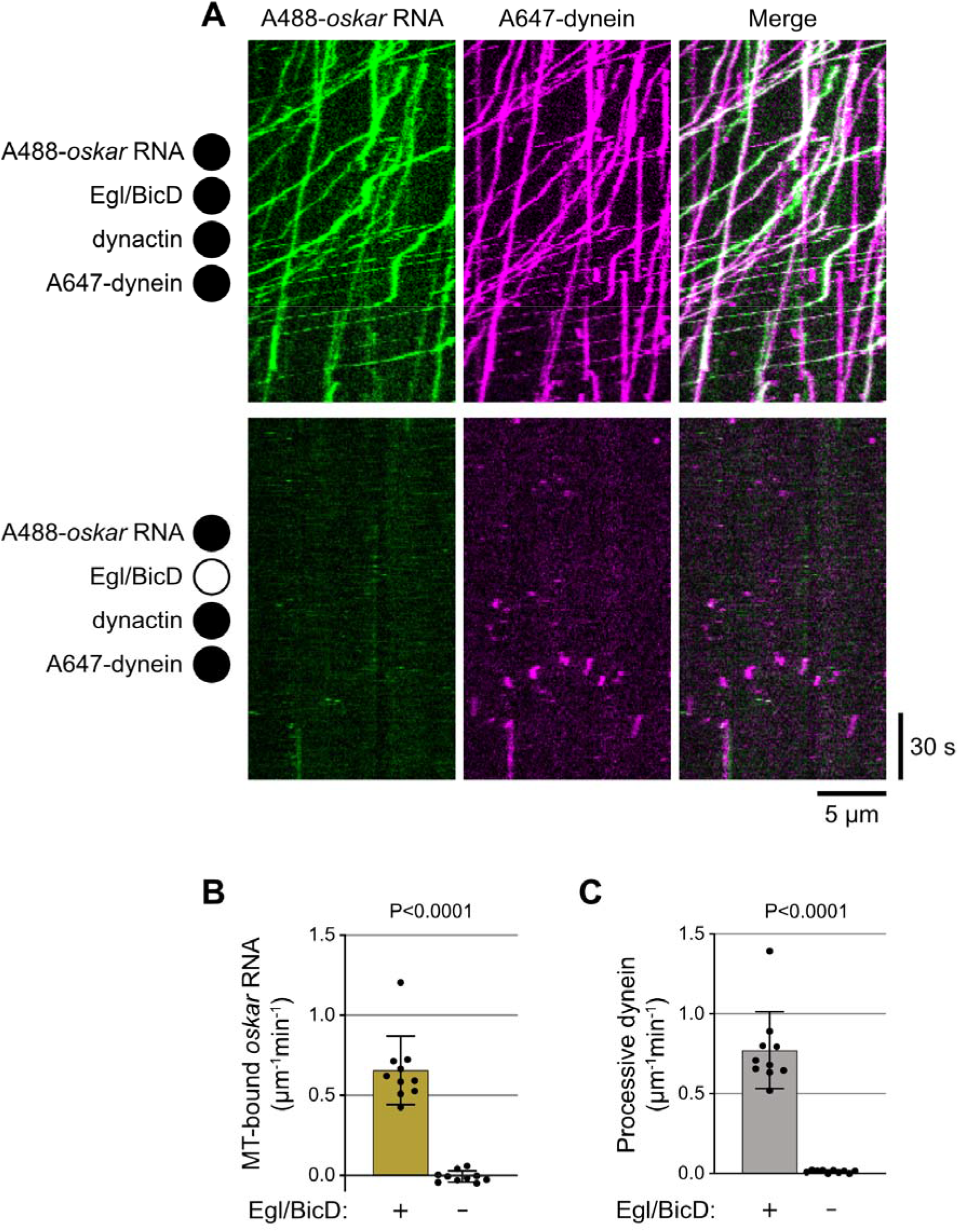
Egl and BicD are required for formation of transport-competent dynein-RNA complexes *in vitro*. (A) Example kymographs (time-distance plots) showing behavior of dynein and *oskar* RNA in the presence and absence of Egl and BicD; in each condition, dynactin is also present but not fluorescently labeled (filled and empty circles represent the presence and absence of indicated proteins, respectively). Egl and BicD were co-expressed and co-purified (see Methods). (B and C) Charts showing the total number of microtubule (MT) binding events for *oskar* RNA (B) and number of processive dynein complexes on microtubules under conditions shown in A. In B, values were corrected for non-specific background binding of *oskar* RNA to the imaging surface as described in Methods. Mean ± SD of values from individual microtubules (represented by black circles) are displayed. Statistical significance and P-values were determined with Mann-Whitney tests.

**Figure S3.**
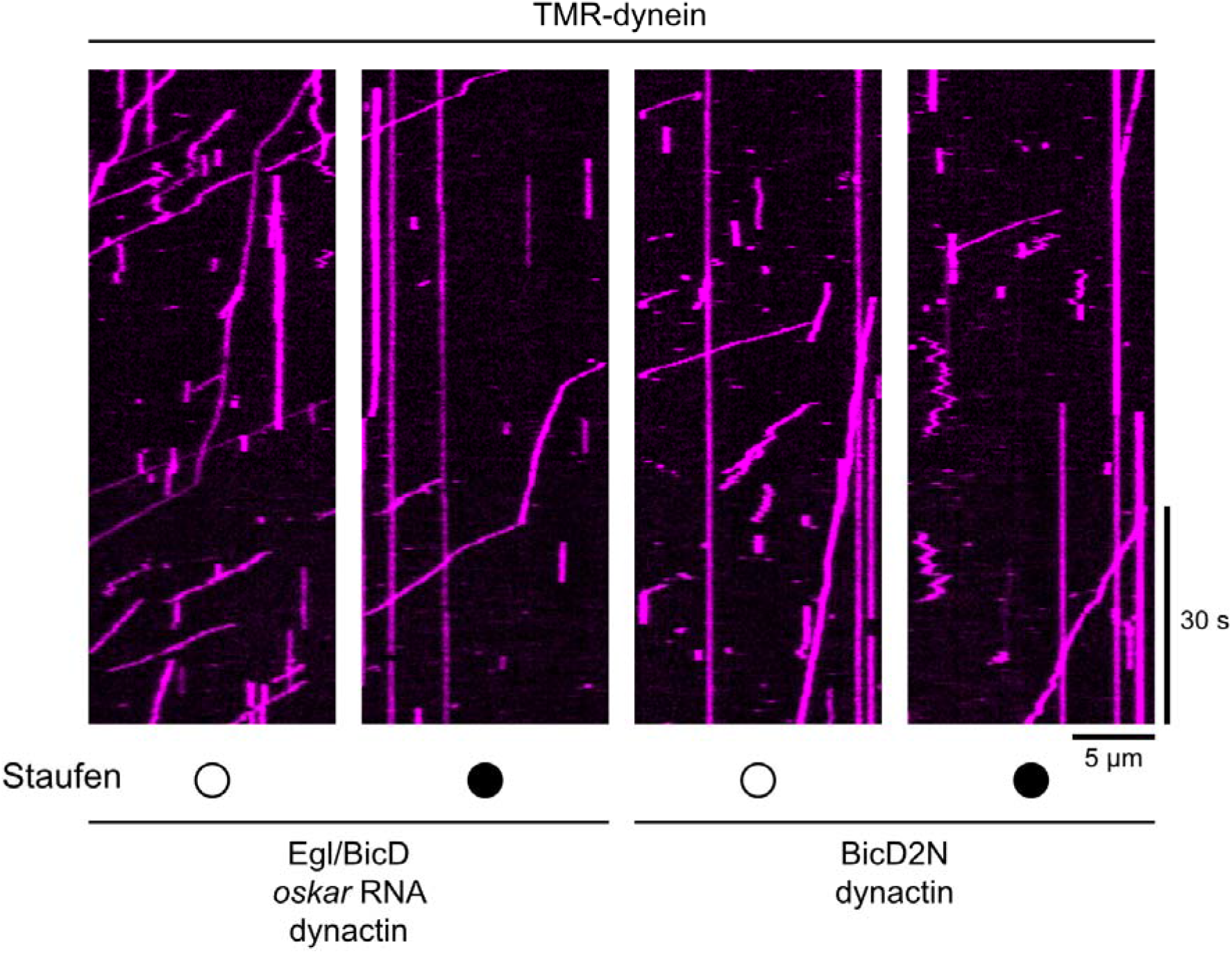
Example kymographs (time-distance plots) showing the behavior of dynein activated by *oskar* RNA, Egl/BicD, and dynactin or by BicD2N and dynactin in the presence (filled circle) and absence (open circle) of Staufen. Quantification of these data is presented in Fig. 2H and I.

**Figure S4:**
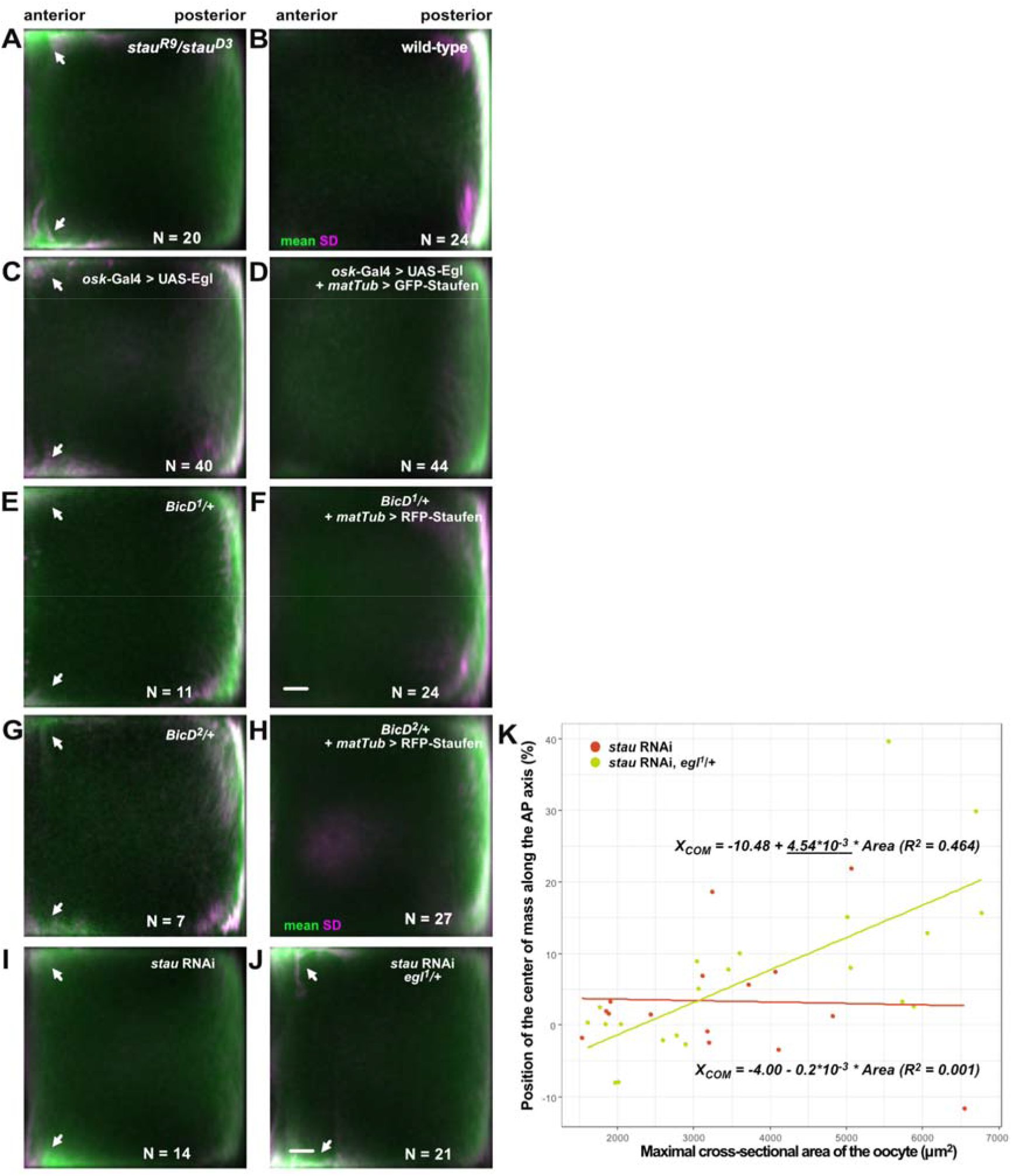
Suppression of *oskar* mislocalization. (A-J) Average distribution of *oskar* mRNA (green) and variability of the distribution (SD, magenta) in stage 9 oocytes of the indicated genotypes. N indicates the number of oocytes analyzed. Scale bar represents 10% of anteroposterior axis length. Anterior is to the left, posterior to the right. Arrowheads indicate the ectopic localization of *oskar* mRNA at the anterolateral corners. (K) Distribution of the observed *oskar* center-of-mass in stage 9 *stau* RNAi oocytes in the presence of two (red) or one (yellow) functional copies of *egl* as a function of oocyte size, used here as a proxy for developmental stage. Solid lines show the best linear fits to the data. The equation and the square of the goodness-of-fit (R^2^) are indicated. Such moderate rescue was expected as *oskar* RNPs entering the oocyte are thought to be associated with Egl. Note that there is no significant linear correlation between oocyte size (developmental stage) and the position of *oskar* RNA center-of-mass in *stau* RNAi (red), indicating an *oskar* mislocalization phenotype. There is a moderate correlation with a significant slope (underlined) when one copy of *egl* is removed (yellow), indicating progressive posterior localization of *oskar* mRNA at stage 9.

**Figure S5:**
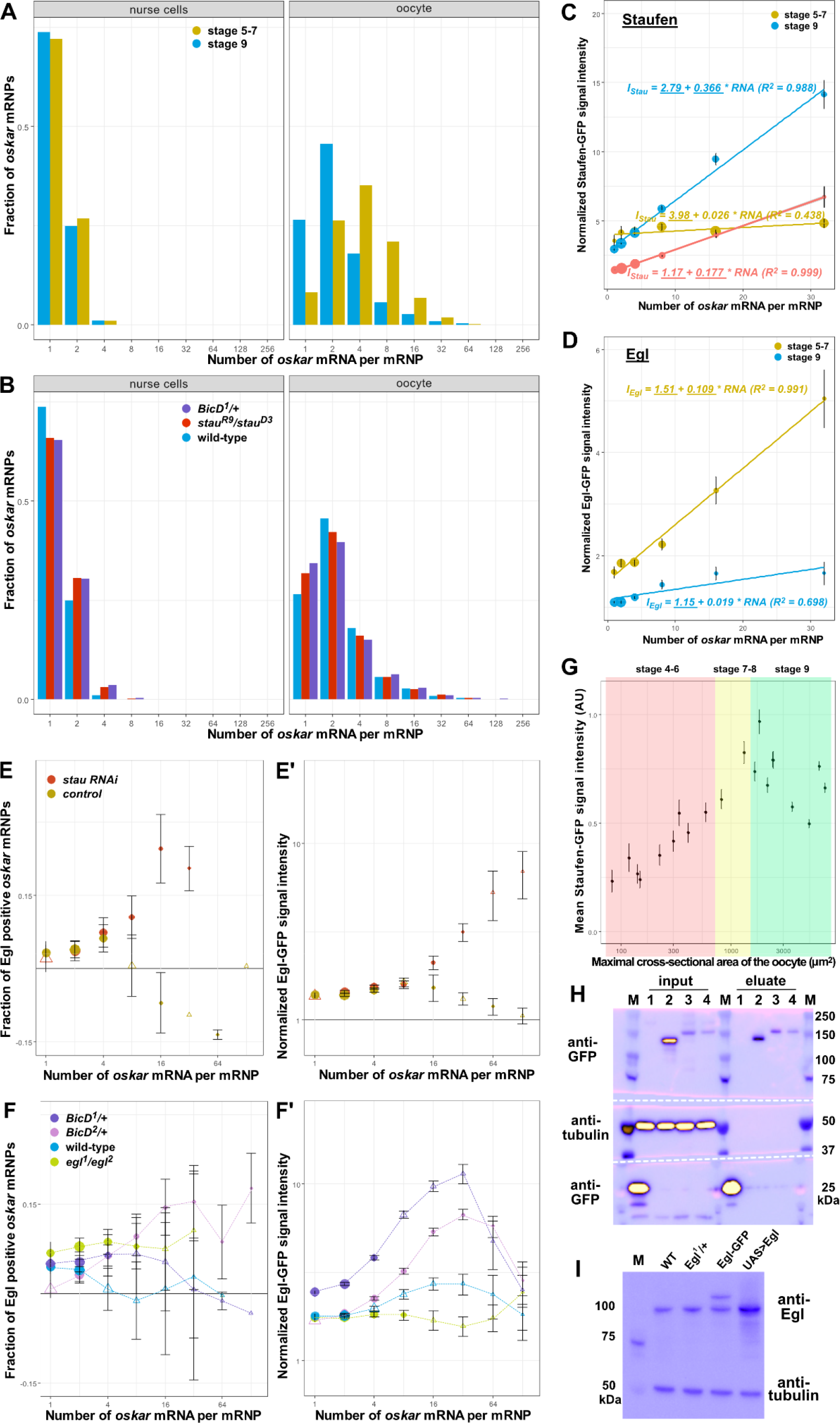
Staufen and Egl association with *oskar* RNPs. (A, B) *oskar* mRNA content distribution in (A) wild-type nurse cells and oocytes at stages 5-7 (yellow) and 9 (blue) of oogenesis, and (B) in *stau* null (red) and dominant *BicD^1^* mutants (purple). In the nurse cells, most *oskar* RNPs contain 1-2 copies of the RNA, consistent with previous reports (Little et al., 2015). *oskar* RNP content increases in the oocyte: at stages 5-7 most RNPs contain 4+ copies of the RNA, which decreases to >2 copies during *oskar* posterior localization (stage 9; Little et al., 2015). (C, D) Normalized Staufen-GFP (B’) or Egl-GFP (D’, E’) signal intensity as a function of *oskar* mRNA content at stages 5-7 (yellow) and stage 9 (blue). (C) Staufen-GFP signal intensity in the complete absence (yellow and blue) and in the presence (red) of endogenous, unlabeled Staufen. Fitted linear models showing the correlation between (C) Staufen-GFP and (D) Egl-GFP signal intensity and *oskar* mRNA copy number as solid lines and equations (top - stages 5-7, bottom - stage 9). Underscored parameters of the models are significantly different from zero (p<0.05). The slopes of the two fitted models are significantly different (p<0.0001, ANOVA). (E-F’) Association of Egl-GFP with *oskar* RNPs in oocytes expressing (E, E’) *stau* RNAi (red), control RNAi (brown), or with (F, F’) *BicD^1^* (purple) or *BicD^2^* (pink) alleles. (E) Note that knock-down of Staufen results in similar retention of Egl on *oskar* RNPs as in the complete absence of Staufen protein (Fig 3E, E’). (D-F’) Egg-chambers expressed a single copy of Egl-GFP in the presence of two endogenous wild-type *egl* alleles, except in the case of the rescued *egl* mutants (F, *egl^1^/egl^2^*, green). Although when unlabelled Egl was absent (*egl^1^/egl^2^*, green), we observed a slightly elevated fraction of Egl positive RNPs, larger RNPs containing 16+ copies of *oskar* mRNA displayed no significant association with Egl (F) and the relative amounts of Egl on *oskar* RNPs were identical to what was observed in the presence of endogenous, unlabelled Egl (F’, blue). (G) Mean signal intensity of Staufen-GFP measured at multiple locations throughout developing oocytes. Size of the oocytes (x-axis) is used as a proxy of developmental time and, along with morphological features, for staging of the oocytes (shaded areas as indicated in the panel). (C-G) Error bars show 95% confidence interval of the mean. (H) Western blot of input lysates and eluates after RNA immunoprecipitation in the presence (lane 3) or the absence (lane 4) of Staufen. Bait proteins - monomeric EGFP (lane 1), GFP-Staufen (lane 2) and Egl-GFP (lanes 3,4) - are detected by anti-GFP antibody. Anti-tubulin staining served to monitor potential contamination of the eluates. (I) Western blot showing endogenous Egl protein detected by anti-Egl antibody in the indicated genotypes. Tubulin was used as a loading control. In H and I, blue and yellow indicate low and high intensity of signal, respectively.

**Figure S6:**
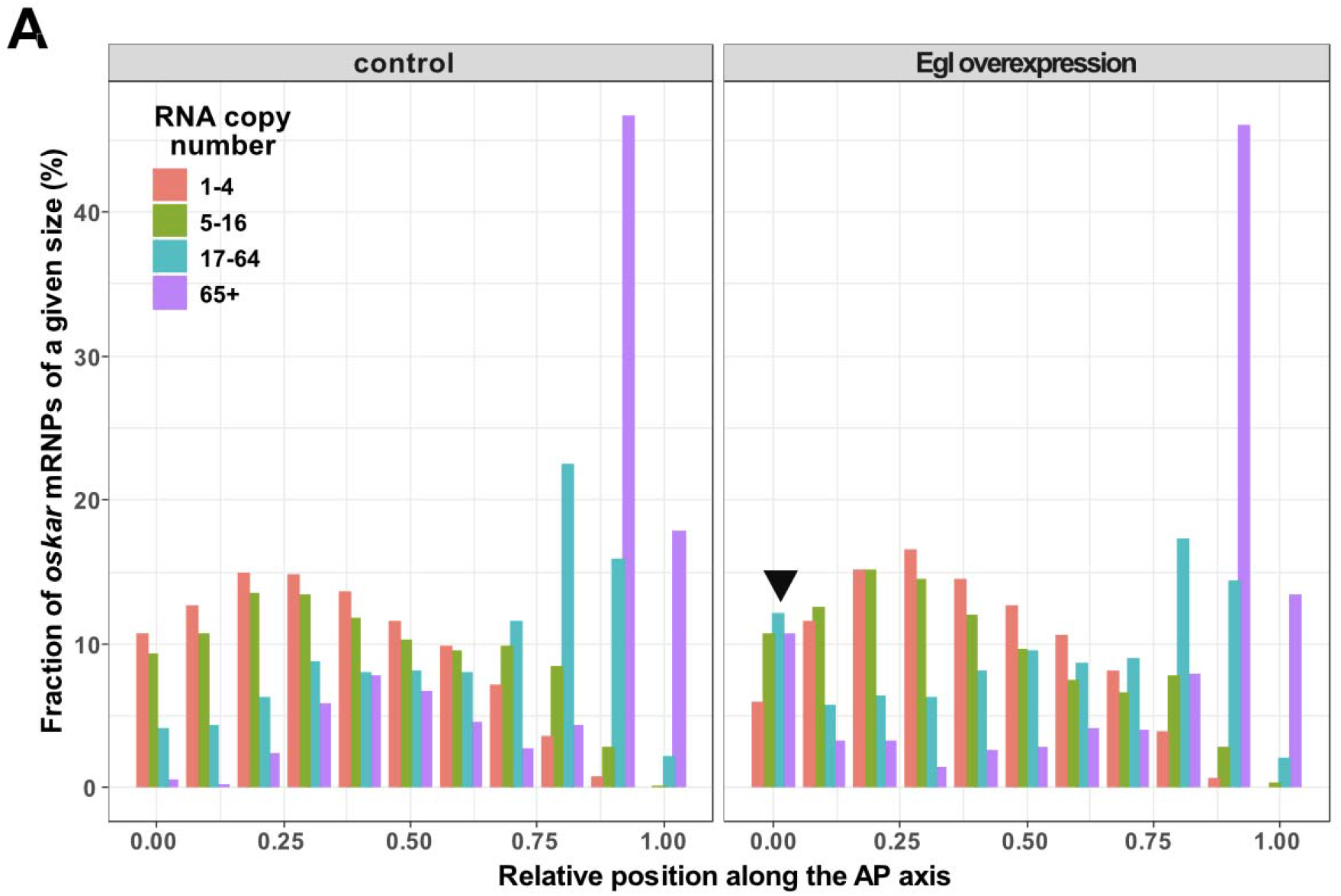
Localization of *oskar* RNPs along the anteroposterior axis. (A) Relative distribution of *oskar* RNPs grouped by RNA content along the anteroposterior axis in wild-type and in Egl overexpressing oocytes.

**Figure S7:**
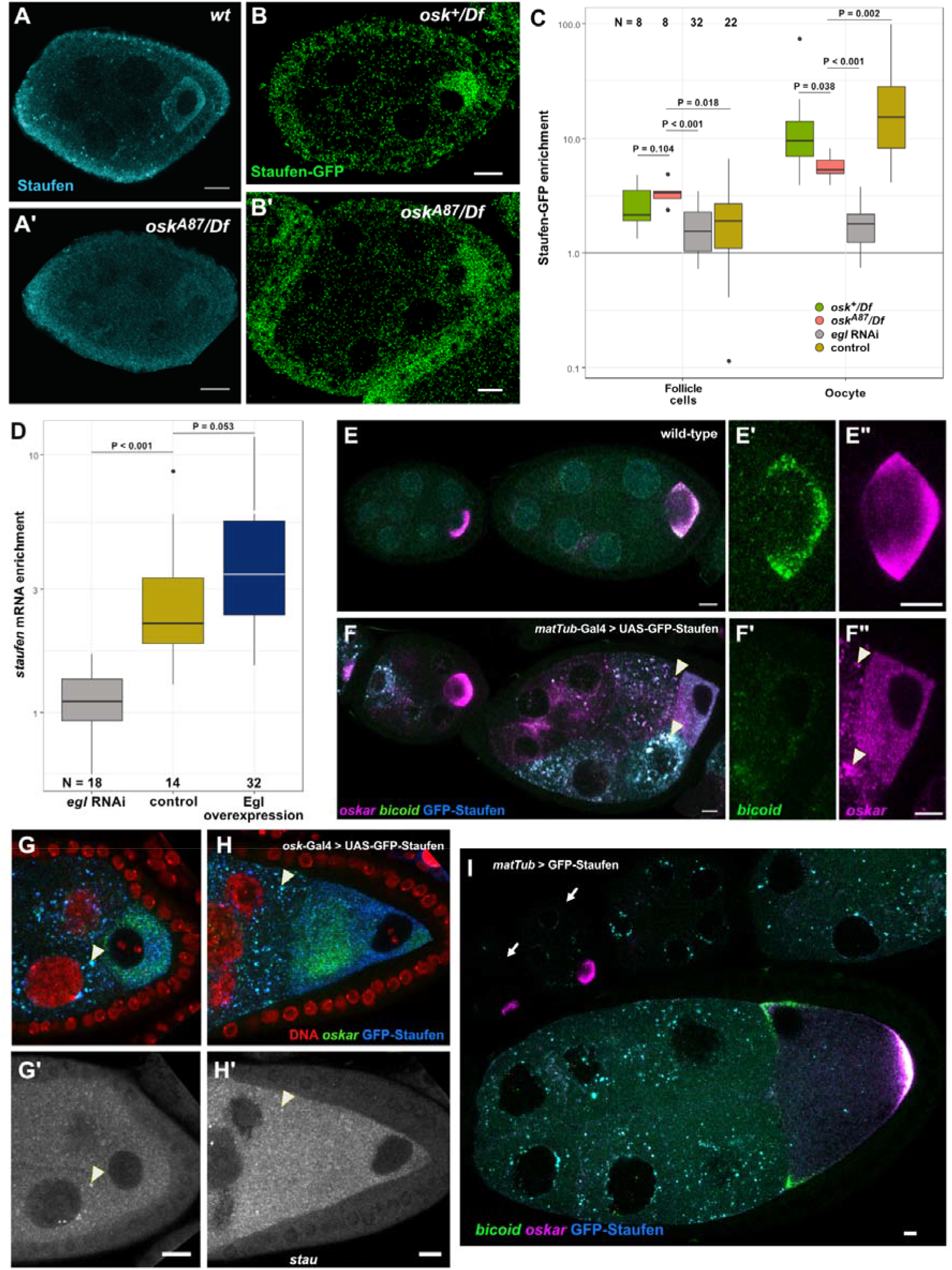
Localization of Staufen protein and *stau* mRNA in the egg-chamber. (A-B’) Staufen protein expression detected by immunofluorescence (blue, A and A’) or by the fluorescent reporter Staufen-GFP (B and B’, green) in early egg-chambers in the presence (A and B) and complete absence of *oskar* mRNA (A’ and B’). Note that lack of *oskar* from the oocyte blocks progression of oogenesis beyond stage 6 (Jenny et al., 2006). (C) Enrichment of Staufen-GFP signal in early oocytes with one (green) or no (red) functional *oskar* alleles expressing *oskar* mRNA, and in oocytes expressing *egl* RNAi (gray) or control RNAi (brown). Enrichment is relative to the sibling nurse cells in the egg-chamber. Enrichment of Staufen-GFP in somatic follicle cells, which do not express the shRNA, serves as a control. P-values of pairwise Student’s t-test are shown. Note that complete lack of *oskar* mRNA, an abundant binding partner of Staufen, has only a moderate effect on Staufen enrichment in the developing oocyte (also observed in A-B’), whereas knock-down of Egl almost completely abolishes Staufen ooplasmic accumulation. (D) Quantification of the enrichment of *stau* mRNA in the oocyte in *egl* RNAi (gray), control RNAi (yellow) and Egl overexpressing oocytes (dark blue). Note that an excess of Egl has a minuscule effect on *stau* RNA accumulation in the oocyte, suggesting that in the wild-type, most of the *stau* mRNA expressed in the nurse cells is transported into the oocyte. (E-F”) Early egg-chambers overexpressing GFP-Staufen (F, blue) under control of the *matTub*-Gal4 driver. Note that in control oocytes of a similar stage (E-E”), and in oocytes expressing low levels of GFP-Staufen (F, left egg-chamber), *oskar* mRNA (E”, F”, magenta) is enriched. Enrichment of *oskar* and *bicoid* (E’, F’, green) mRNAs is greatly reduced in oocytes expressing high levels of GFP-Staufen (F, right egg-chamber). This phenotype is reproducibly observed in >30 egg-chambers derived from three separate crosses. (G-H’) Egg-chambers in early and mid-oogenesis overexpressing GFP-Staufen (blue) under control of the *oskar*-Gal4 driver from the beginning of oogenesis. The vast majority of such egg-chambers fail to develop beyond stage 6, likely as a consequence of greatly reduced ooplasmic accumulation of *oskar* mRNA (green), which appears to be trapped in the nurse cells in large aggregates associated with GFP-Staufen (some examples are highlighted by arrowheads in panels F-H’). Such aggregates of endogenous Staufen are not observed in wild-type egg-chambers (E-E”). Similarly, no accumulation of *stau* mRNA in the oocyte is observed in these *oskar-Gal4>UAS-GFP-Staufen* oocytes (G’ and H’), where *stau* mRNA levels are uniformly high in the germline (compare the signal in the follicle cell layer to that in the nurse cells and in the oocyte; see Figure 7). (H, H’) In the oocytes occasionally escaping early developmental arrest, we invariably observed failure in nuclear migration from the posterior to the anterior, reflecting a defect in repolarization of the oocyte microtubule network (Januschke et al., 2006). Consequently, *oskar* mRNA remains in the center of these oocytes, which - although they complete oogenesis - fail to result in viable progeny. These phenotypes are reproducibly observed in >30 egg-chambers derived from three separate crosses. (I) Expression of GFP-Staufen under the control of maternal tubulin promoter. GFP-Staufen (cyan) can hardly be detected in early egg chambers (highlighted by arrows), and the forming aggregates remain associated with the nurse cell nuclei until mid-oogenesis. Scale bars represent 5 μm.

## REFERENCES

Abouward, R., and G. Schiavo. 2021. Walking the line: mechanisms underlying directional mRNA transport and localisation in neurons and beyond. Cell. Mol. Life Sci. 78:2665–2681. doi:10.1007/s00018-020-03724-3.

Amrute-Nayak, M., and S.L. Bullock. 2012. Single-molecule assays reveal that RNA localization signals regulate dynein-dynactin copy number on individual transcript cargoes. Nat. Cell Biol. 14:416–423. doi:10.1038/ncb2446.

Bauer, K.E., I. Segura, I. Gaspar, V. Scheuss, C. Illig, G. Ammer, S. Hutten, E. Basyuk, S.M. Fernández-Moya, J. Ehses, E. Bertrand, and M.A. Kiebler. 2019. Live cell imaging reveals 3’-UTR dependent mRNA sorting to synapses. Nat. Commun. 10:3178. doi:10.1038/s41467-019-11123-x.

Baumann, S., T. Pohlmann, M. Jungbluth, A. Brachmann, and M. Feldbrügge. 2012. Kinesin-3 and dynein mediate microtubule-dependent co-transport of mRNPs and endosomes. J. Cell Sci. 125:2740–2752. doi:10.1242/jcs.101212.

Benaglia, T., D. Chauveau, D.R. Hunter, and D. Young. 2009. mixtoolsL: An R package for Analyzing Finite Mixture Models. J. Stat. Softw. 32. doi:10.18637/jss.v032.i06.

Berleth, T., M. Burri, G. Thoma, D. Bopp, S. Richstein, G. Frigerio, M. Noll, and C. Nüsslein-Volhard. 1988. The role of localization of bicoid RNA in organizing the anterior pattern of the Drosophila embryo. EMBO J. 7:1749–1756. doi:10.1002/j.1460-2075.1988.tb03004.x.

Brendza, R.P., L.R. Serbus, J.B. Duffy, and W.M. Saxton. 2000. A function for kinesin I in the posterior transport of oskar mRNA and Staufen protein. Science. 289:2120–2122. doi:10.1126/science.289.5487.2120.

Bullock, S.L., A. Nicol, S.P. Gross, and D. Zicha. 2006. Guidance of bidirectional motor complexes by mRNA cargoes through control of dynein number and activity. Curr. Biol. 16:1447–1452. doi:10.1016/j.cub.2006.05.055.

Clark, A., C. Meignin, and I. Davis. 2007. A Dynein-dependent shortcut rapidly delivers axis determination transcripts into the Drosophila oocyte. Development. 134:1955–1965. doi:10.1242/dev.02832.

Dienstbier, M., F. Boehl, X. Li, and S.L. Bullock. 2009. Egalitarian is a selective RNA-binding protein linking mRNA localization signals to the dynein motor. Genes Dev. 23:1546–1558. doi:10.1101/gad.531009.

Dix, C.I., H.C. Soundararajan, N.S. Dzhindzhev, F. Begum, B. Suter, H. Ohkura, E. Stephens, and S.L. Bullock. 2013. Lissencephaly-1 promotes the recruitment of dynein and dynactin to transported mRNAs. J. Cell Biol. 202:479–494. doi:10.1083/jcb.201211052.

Edelstein, A.D., M.A. Tsuchida, N. Amodaj, H. Pinkard, R.D. Vale, and N. Stuurman. 2014. Advanced methods of microscope control using μManager software. J. Biol. Methods. 1. doi:10.14440/jbm.2014.36.

van Eeden, F.J., I.M. Palacios, M. Petronczki, M.J. Weston, and D. St Johnston. 2001. Barentsz is essential for the posterior localization of oskar mRNA and colocalizes with it to the posterior pole. J. Cell Biol. 154:511–523. doi:10.1083/jcb.200105056.

Ephrussi, A., L.K. Dickinson, and R. Lehmann. 1991. Oskar organizes the germ plasm and directs localization of the posterior determinant nanos. Cell. 66:37–50. doi:10.1016/0092-8674(91)90137-n.

Ephrussi, A., and R. Lehmann. 1992. Induction of germ cell formation by oskar. Nature. 358:387–392. doi:10.1038/358387a0.

Ferrandon, D., L. Elphick, C. Nüsslein-Volhard, and D. St Johnston. 1994. Staufen protein associates with the 3’UTR of bicoid mRNA to form particles that move in a microtubule-dependent manner. Cell. 79:1221–1232. doi:10.1016/0092-8674(94)90013-2.

Gagnon, J.A., J.A. Kreiling, E.A. Powrie, T.R. Wood, and K.L. Mowry. 2013. Directional transport is mediated by a Dynein-dependent step in an RNA localization pathway. PLoS Biol. 11:e1001551. doi:10.1371/journal.pbio.1001551.

Gaspar, I., and A. Ephrussi. 2017. Ex vivo Ooplasmic Extract from Developing Drosophila Oocytes for Quantitative TIRF Microscopy Analysis. Bio Protoc. 7. doi:10.21769/BioProtoc.2380.

Gáspár, I., V. Sysoev, A. Komissarov, and A. Ephrussi. 2017. An RNA-binding atypical tropomyosin recruits kinesin-1 dynamically to oskar mRNPs. EMBO J. 36:319–333. doi:10.15252/embj.201696038.

Gaspar, I., F. Wippich, and A. Ephrussi. 2017. Enzymatic production of single-molecule FISH and RNA capture probes. RNA. 23:1582–1591. doi:10.1261/rna.061184.117.

Gaspar, I., Y.V. Yu, S.L. Cotton, D.-H. Kim, A. Ephrussi, and M.A. Welte. 2014. Klar ensures thermal robustness of oskar localization by restraining RNP motility. J. Cell Biol. 206:199–215. doi:10.1083/jcb.201310010.

Gaspar, I. 2011. Microtubule-based motor-mediated mRNA localization in Drosophila oocytes and embryos. Biochem. Soc. Trans. 39:1197–1201. doi:10.1042/BST0391197.

Ghosh, S., V. Marchand, I. Gáspár, and A. Ephrussi. 2012. Control of RNP motility and localization by a splicing-dependent structure in oskar mRNA. Nat. Struct. Mol. Biol. 19:441–449. doi:10.1038/nsmb.2257.

Glock, C., M. Heumüller, and E.M. Schuman. 2017. mRNA transport & local translation in neurons. Curr. Opin. Neurobiol. 45:169–177. doi:10.1016/j.conb.2017.05.005.

Gratz, S.J., A.M. Cummings, J.N. Nguyen, D.C. Hamm, L.K. Donohue, M.M. Harrison, J. Wildonger, and K.M. O’Connor-Giles. 2013. Genome engineering of Drosophila with the CRISPR RNA-guided Cas9 nuclease. Genetics. 194:1029–1035. doi:10.1534/genetics.113.152710.

Hachet, O., and A. Ephrussi. 2001. Drosophila Y14 shuttles to the posterior of the oocyte and is required for oskar mRNA transport. Curr. Biol. 11:1666–1674. doi:10.1016/s0960-9822(01)00508-5.

Hachet, O., and A. Ephrussi. 2004. Splicing of oskar RNA in the nucleus is coupled to its cytoplasmic localization. Nature. 428:959–963. doi:10.1038/nature02521.

Hancock, W.O. 2014. Bidirectional cargo transport: moving beyond tug of war. Nat. Rev. Mol. Cell Biol. 15:615–628. doi:10.1038/nrm3853.

Heber, S., I. Gáspár, J.-N. Tants, J. Günther, S.M.F. Moya, R. Janowski, A. Ephrussi, M. Sattler, and D. Niessing. 2019. Staufen2-mediated RNA recognition and localization requires combinatorial action of multiple domains. Nat. Commun. 10:1659. doi:10.1038/s41467-019-09655-3.

Heraud-Farlow, J.E., T. Sharangdhar, X. Li, P. Pfeifer, S. Tauber, D. Orozco, A. Hörmann, S. Thomas, A. Bakosova, A.R. Farlow, D. Edbauer, H.D. Lipshitz, Q.D. Morris, M. Bilban, M. Doyle, and M.A. Kiebler. 2013. Staufen2 regulates neuronal target RNAs. Cell Rep. 5:1511–1518. doi:10.1016/j.celrep.2013.11.039.

Hoang, H.T., M.A. Schlager, A.P. Carter, and S.L. Bullock. 2017. DYNC1H1 mutations associated with neurological diseases compromise processivity of dynein-dynactin-cargo adaptor complexes. Proc Natl Acad Sci USA. 114:E1597–E1606. doi:10.1073/pnas.1620141114.

Hoogenraad, C.C., and A. Akhmanova. 2016. Bicaudal D family of motor adaptors: linking dynein motility to cargo binding. Trends Cell Biol. 26:327–340. doi:10.1016/j.tcb.2016.01.001.

Hoogenraad, C.C., P. Wulf, N. Schiefermeier, T. Stepanova, N. Galjart, J.V. Small, F. Grosveld, C.I. de Zeeuw, and A. Akhmanova. 2003. Bicaudal D induces selective dynein-mediated microtubule minus end-directed transport. EMBO J. 22:6004–6015. doi:10.1093/emboj/cdg592.

Jambor, H., S. Mueller, S.L. Bullock, and A. Ephrussi. 2014. A stem-loop structure directs oskar mRNA to microtubule minus ends. RNA. 20:429–439. doi:10.1261/rna.041566.113.

Januschke, J., L. Gervais, S. Dass, J.A. Kaltschmidt, H. Lopez-Schier, D. St Johnston, A.H. Brand, S. Roth, and A. Guichet. 2002. Polar transport in the Drosophila oocyte requires Dynein and Kinesin I cooperation. Curr. Biol. 12:1971–1981. doi:10.1016/S0960-9822(02)01302-7.

Januschke, J., L. Gervais, L. Gillet, G. Keryer, M. Bornens, and A. Guichet. 2006. The centrosome-nucleus complex and microtubule organization in the Drosophila oocyte. Development. 133:129–139. doi:10.1242/dev.02179.

Jenny, A., O. Hachet, P. Závorszky, A. Cyrklaff, M.D.J. Weston, D.S. Johnston, M. Erdélyi, and A. Ephrussi. 2006. A translation-independent role of oskar RNA in early Drosophila oogenesis. Development. 133:2827–2833. doi:10.1242/dev.02456.

Kim-Ha, J., J.L. Smith, and P.M. Macdonald. 1991. oskar mRNA is localized to the posterior pole of the Drosophila oocyte. Cell. 66:23–35. doi:10.1016/0092-8674(91)90136-m.

Lasko, P. 2012. mRNA localization and translational control in Drosophila oogenesis. Cold Spring Harb. Perspect. Biol. 4. doi:10.1101/cshperspect.a012294.

Laver, J.D., X. Li, K. Ancevicius, J.T. Westwood, C.A. Smibert, Q.D. Morris, and H.D. Lipshitz. 2013. Genome-wide analysis of Staufen-associated mRNAs identifies secondary structures that confer target specificity. Nucleic Acids Res. 41:9438–9460. doi:10.1093/nar/gkt702.

Lenth, R.V. 2016. Least-Squares Means: the R package lsmeans. J. Stat. Softw. 69:1–33. doi:10.18637/jss.v069.i01.

Little, S.C., K.S. Sinsimer, J.J. Lee, E.F. Wieschaus, and E.R. Gavis. 2015. Independent and coordinate trafficking of single Drosophila germ plasm mRNAs. Nat. Cell Biol. 17:558–568. doi:10.1038/ncb3143.

Liu, Y., H.K. Salter, A.N. Holding, C.M. Johnson, E. Stephens, P.J. Lukavsky, J. Walshaw, and S.L. Bullock. 2013. Bicaudal-D uses a parallel, homodimeric coiled coil with heterotypic registry to coordinate recruitment of cargos to dynein. Genes Dev. 27:1233–1246. doi:10.1101/gad.212381.112.

Mach, J.M., and R. Lehmann. 1997. An Egalitarian-BicaudalD complex is essential for oocyte specification and axis determination in Drosophila. Genes Dev. 11:423–435. doi:10.1101/gad.11.4.423.

Marchand, V., I. Gaspar, and A. Ephrussi. 2012. An intracellular transmission control protocol: assembly and transport of ribonucleoprotein complexes. Curr. Opin. Cell Biol. 24:202–210. doi:10.1016/j.ceb.2011.12.014.

Martin, K.C., and A. Ephrussi. 2009. mRNA localization: gene expression in the spatial dimension. Cell. 136:719–730. doi:10.1016/j.cell.2009.01.044.

McClintock, M.A., C.I. Dix, C.M. Johnson, S.H. McLaughlin, R.J. Maizels, H.T. Hoang, and S.L. Bullock. 2018. RNA-directed activation of cytoplasmic dynein-1 in reconstituted transport RNPs. eLife. 7. doi:10.7554/eLife.36312.

McKenney, R.J., W. Huynh, M.E. Tanenbaum, G. Bhabha, and R.D. Vale. 2014. Activation of cytoplasmic dynein motility by dynactin-cargo adapter complexes. Science. 345:337–341. doi:10.1126/science.1254198.

McKenney, R.J., M. Vershinin, A. Kunwar, R.B. Vallee, and S.P. Gross. 2010. LIS1 and NudE induce a persistent dynein force-producing state. Cell. 141:304–314. doi:10.1016/j.cell.2010.02.035.

Micklem, D.R., J. Adams, S. Grünert, and D. St Johnston. 2000. Distinct roles of two conserved staufen domains in oskar mRNA localization and translation. EMBO J. 19:1366–1377. doi:10.1093/emboj/19.6.1366.

Mofatteh, M., and S.L. Bullock. 2017. SnapShot: Subcellular mRNA Localization. Cell. 169:178–178.e1. doi:10.1016/j.cell.2017.03.004.

Mohler, J., and E.F. Wieschaus. 1986. Dominant maternal-effect mutations of Drosophila melanogaster causing the production of double-abdomen embryos. Genetics. 112:803–822. doi:10.1093/genetics/112.4.803.

Mohr, S., A. Kenny, S.T.Y. Lam, M.B. Morgan, C.A. Smibert, H.D. Lipshitz, and P.M. Macdonald. 2021. Opposing roles for Egalitarian and Staufen in transport, anchoring and localization of oskar mRNA in the Drosophila oocyte. PLoS Genet. 17:e1009500. doi:10.1371/journal.pgen.1009500.

Mohr, S.E., S.T. Dillon, and R.E. Boswell. 2001. The RNA-binding protein Tsunagi interacts with Mago Nashi to establish polarity and localize oskar mRNA during Drosophila oogenesis. Genes Dev. 15:2886–2899. doi:10.1101/gad.927001.

Navarro, C., H. Puthalakath, J.M. Adams, A. Strasser, and R. Lehmann. 2004. Egalitarian binds dynein light chain to establish oocyte polarity and maintain oocyte fate. Nat. Cell Biol. 6:427–435. doi:10.1038/ncb1122.

Neuman-Silberberg, F.S., and T. Schüpbach. 1993. The Drosophila dorsoventral patterning gene gurken produces a dorsally localized RNA and encodes a TGFα-like protein. Cell. 75:165–174. doi:10.1016/S0092-8674(05)80093-5.

Newmark, P.A., and R.E. Boswell. 1994. The mago nashi locus encodes an essential product required for germ plasm assembly in Drosophila. Development. 120:1303–1313.

O’Connor-Giles, Wildonger, and Harrison. 2014. Generating targeting chiRNAs. FlyCRISPR Web Site.

Palacios, I.M., D. Gatfield, D. St Johnston, and E. Izaurralde. 2004. An eIF4AIII-containing complex required for mRNA localization and nonsense-mediated mRNA decay. Nature. 427:753–757. doi:10.1038/nature02351.

Palacios, I.M., and D. St Johnston. 2002. Kinesin light chain-independent function of the Kinesin heavy chain in cytoplasmic streaming and posterior localisation in the Drosophila oocyte. Development. 129:5473–5485. doi:10.1242/dev.00119.

Parton, R.M., R.S. Hamilton, G. Ball, L. Yang, C.F. Cullen, W. Lu, H. Ohkura, and I. Davis. 2011. A PAR-1-dependent orientation gradient of dynamic microtubules directs posterior cargo transport in the Drosophila oocyte. J. Cell Biol. 194:121–135. doi:10.1083/jcb.201103160.

R Core Team. 2014. R: A language and environment for statistical computing. R Foundation for StatisticalComputing.

Sanghavi, P., S. Laxani, X. Li, S.L. Bullock, and G.B. Gonsalvez. 2013. Dynein associates with oskar mRNPs and is required for their efficient net plus-end localization in Drosophila oocytes. PLoS ONE. 8:e80605. doi:10.1371/journal.pone.0080605.

Sanghavi, P., G. Liu, R. Veeranan-Karmegam, C. Navarro, and G.B. Gonsalvez. 2016. Multiple roles for Egalitarian in polarization of the Drosophila egg chamber. Genetics. 203:415–432. doi:10.1534/genetics.115.184622.

Schindelin, J., I. Arganda-Carreras, E. Frise, V. Kaynig, M. Longair, T. Pietzsch, S. Preibisch, C. Rueden, S. Saalfeld, B. Schmid, J.-Y. Tinevez, D.J. White, V. Hartenstein, K. Eliceiri, P. Tomancak, and A. Cardona. 2012. Fiji: an open-source platform for biological-image analysis. Nat. Methods. 9:676–682. doi:10.1038/nmeth.2019.

Schlager, M.A., H.T. Hoang, L. Urnavicius, S.L. Bullock, and A.P. Carter. 2014. In vitro reconstitution of a highly processive recombinant human dynein complex. EMBO J. 33:1855–1868. doi:10.15252/embj.201488792.

Schneider, C.A., W.S. Rasband, and K.W. Eliceiri. 2012. NIH Image to ImageJ: 25 years of image analysis. Nat. Methods. 9:671–675. doi:10.1038/nmeth.2089.

Schuldt, A.J., J.H. Adams, C.M. Davidson, D.R. Micklem, J. Haseloff, D. St Johnston, and A.H. Brand. 1998. Miranda mediates asymmetric protein and RNA localization in the developing nervous system. Genes Dev. 12:1847–1857. doi:10.1101/gad.12.12.1847.

Sladewski, T.E., N. Billington, M.Y. Ali, C.S. Bookwalter, H. Lu, E.B. Krementsova, T.A. Schroer, and K.M. Trybus. 2018. Recruitment of two dyneins to an mRNA-dependent Bicaudal D transport complex. eLife. 7. doi:10.7554/eLife.36306.

Soundararajan, H.C., and S.L. Bullock. 2014. The influence of dynein processivity control, MAPs, and microtubule ends on directional movement of a localising mRNA. eLife. 3:e01596. doi:10.7554/eLife.01596.

St Johnston, D., D. Beuchle, and C. Nüsslein-Volhard. 1991. Staufen, a gene required to localize maternal RNAs in the Drosophila egg. Cell. 66:51–63. doi:10.1016/0092-8674(91)90138-o.

St Johnston, D., N.H. Brown, J.G. Gall, and M. Jantsch. 1992. A conserved double-stranded RNA-binding domain. Proc Natl Acad Sci USA. 89:10979–10983. doi:10.1073/pnas.89.22.10979.

St Johnston, D. 2005. Moving messages: the intracellular localization of mRNAs. Nat. Rev. Mol. Cell Biol. 6:363–375. doi:10.1038/nrm1643.

Theurkauf, W.E., B.M. Alberts, Y.N. Jan, and T.A. Jongens. 1993. A central role for microtubules in the differentiation of Drosophila oocytes. Development. 118:1169–1180. doi:10.1242/dev.118.4.1169.

Turner-Bridger, B., M. Jakobs, L. Muresan, H.H.-W. Wong, K. Franze, W.A. Harris, and C.E. Holt. 2018. Single-molecule analysis of endogenous β-actin mRNA trafficking reveals a mechanism for compartmentalized mRNA localization in axons. Proc Natl Acad Sci USA. 115:E9697–E9706. doi:10.1073/pnas.1806189115.

Urnavicius, L., K. Zhang, A.G. Diamant, C. Motz, M.A. Schlager, M. Yu, N.A. Patel, C.V. Robinson, and A.P. Carter. 2015. The structure of the dynactin complex and its interaction with dynein. Science. 347:1441–1446. doi:10.1126/science.aaa4080.

Vazquez-Pianzola, P., B. Schaller, M. Colombo, D. Beuchle, S. Neuenschwander, A. Marcil, R. Bruggmann, and B. Suter. 2017. The mRNA transportome of the BicD/Egl transport machinery. RNA Biol. 14:73–89. doi:10.1080/15476286.2016.1251542.

Wickham, H. 2016. ggplot2 - Elegant Graphics for Data Analysis. Springer-Verlag New York, New York, NY.

Williams, L.S., S. Ganguly, P. Loiseau, B.F. Ng, and I.M. Palacios. 2014. The auto-inhibitory domain and ATP-independent microtubule-binding region of Kinesin heavy chain are major functional domains for transport in the Drosophila germline. Development. 141:176–186. doi:10.1242/dev.097592.

Yoon, Y.J., and K.L. Mowry. 2004. Xenopus Staufen is a component of a ribonucleoprotein complex containing Vg1 RNA and kinesin. Development. 131:3035–3045. doi:10.1242/dev.01170.

Zappulo, A., D. van den Bruck, C. Ciolli Mattioli, V. Franke, K. Imami, E. McShane, M. Moreno-Estelles, L. Calviello, A. Filipchyk, E. Peguero-Sanchez, T. Müller, A. Woehler, C. Birchmeier, E. Merino, N. Rajewsky, U. Ohler, E.O. Mazzoni, M. Selbach, A. Akalin, and M. Chekulaeva. 2017. RNA localization is a key determinant of neurite-enriched proteome. Nat. Commun. 8:583. doi:10.1038/s41467-017-00690-6.

Zimyanin, V.L., K. Belaya, J. Pecreaux, M.J. Gilchrist, A. Clark, I. Davis, and D. St Johnston. 2008. In vivo imaging of oskar mRNA transport reveals the mechanism of posterior localization. Cell. 134:843–853. doi:10.1016/j.cell.2008.06.053.

